# Evolution of membrane rotary ATP synthases: A trade-off between the net productivity and mechanistic stability?

**DOI:** 10.64898/2026.10.01.756066

**Authors:** A.Y. Mulkidjanian, M.V. Kuchevskiy, D.V. Dibrova

## Abstract

We scrutinized the evolution of rotary membrane ATPases/ATP synthases by combining phylogenomic and comparative structural analyses. Regarding F-type ATPases, we found that organisms with more subunits in the membrane-embedded "proteolipid" c-rings tend to have shorter or absent δ-subunits. Organisms with such F-ATPases branch earlier in the phylogenetic tree of catalytic β-subunits. Additionally, we discovered that archaeal-type A-ATPases form at least three distinct branches, including both archaeal and bacterial A-ATPases. This finding suggests that A-ATPase genes have undergone several lateral transfers from archaea to bacteria. We posit that the common ancestor of rotary membrane ATPases possessed six catalytic subunits and over 20 subunits in their Na⁺-transporting c-rings. This ancestor likely had two ion-translocating peripheral stalks and was optimized for low membrane potentials. The loss of half the catalytic subunits, which occurred separately in bacterial and archaeal lineages, enabled the number of c-subunits to be halved without losing efficiency. As cell membranes gained the ability to hold increasingly high membrane voltages, the loss of one of the ion-translocating peripheral stalks, separately in A- and F-ATPases, doubled their efficiency. The mechanism of this loss differed between the two types of ATPases; in F-ATPases, the gain in efficiency came at the cost of mechanistic stability. Thus, the large-scale lateral gene transfer of A-ATPases to bacteria may have been caused by the lower mechanical stability of prokaryotic F-ATPases. The concomitant acquisition of novel subunits resulted in the now-typical Na⁺- and H⁺-transporting A- and F-ATPases geared toward high membrane potentials. Overall, rotary ATP synthases evolved to produce more net ATP.

## 1. Introduction

Rotary ATP synthases produce the lion’s share of ATP in a cell. They couple the synthesis or hydrolysis of ATP in their cytoplasm-protruding, mushroom-like catalytic parts with the sequential translocation of protons or Na^+^ ions across the membrane [1–11]. In the synthesis mode, the transmembrane difference in the electrochemical potentials of protons (Δμ̅H^+^) or Na^+^ ions ( Δμ̅Na^+^) drives the ATP synthesis from ADP and inorganic phosphate. In the hydrolysis mode, the free energy of ATP can drive the formation of Δμ̅H^+^ or Δμ̅ Na^+^ [12,13]. Hereinafter, we will use the term "membrane potential" to refer to both.

All free-living organisms have rotary ATPases, so they are believed to have existed in their common ancestor, known as the Last Universal Cellular (or Common) Ancestor (LUCA) [14]. After the tree of life split into bacterial, archaeal, and eukaryotic branches, the ancestral rotary ATPases evolved into distinguishable bacterial (F-type), archaeal (A-type), and eukaryotic (V-type) versions (Fig. 1A).

**Fig. 1.**
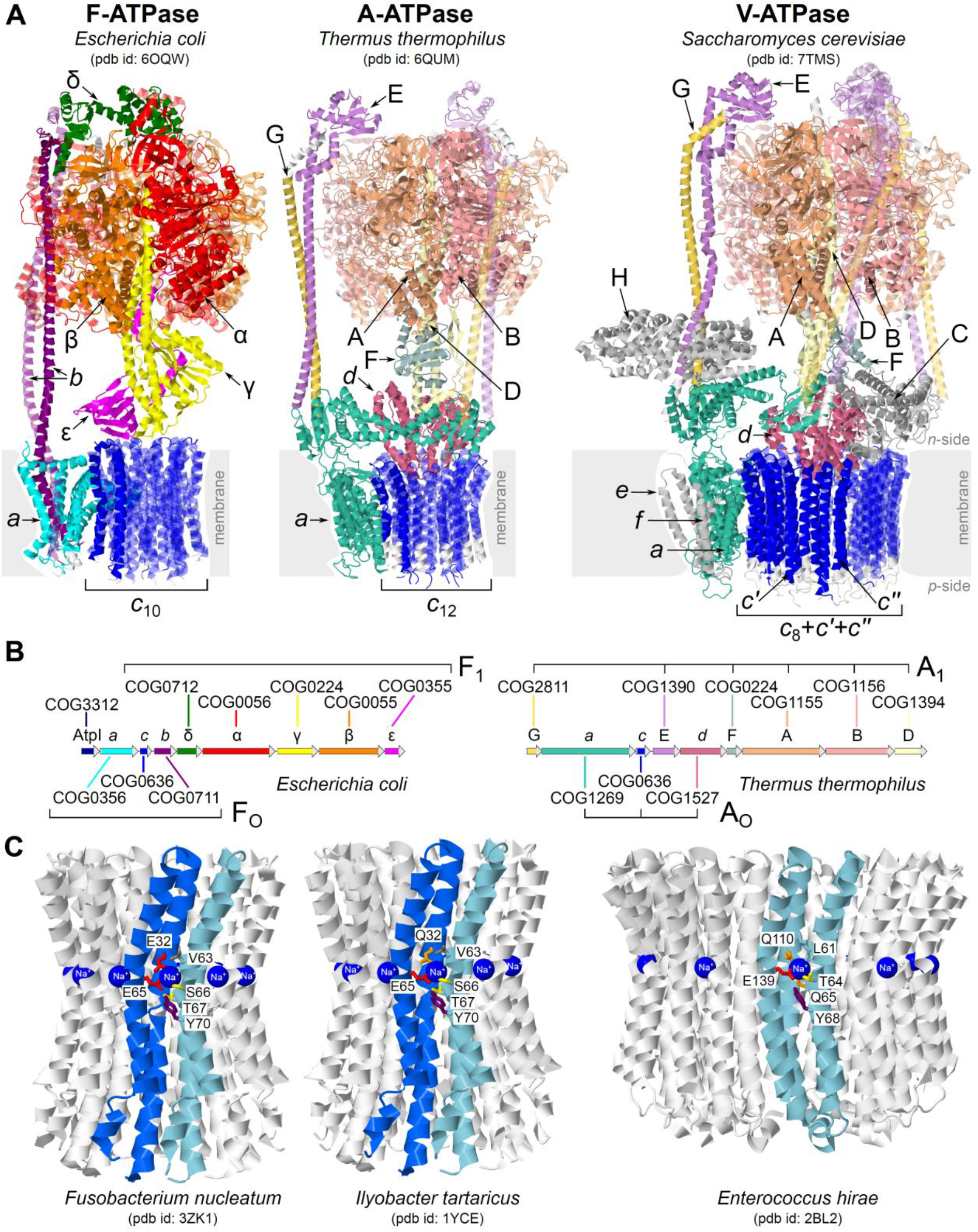
Rotary membrane ATPases. The color scheme used for the subunits and genes of rotary ATPases in panels (**A**) and (**B**) is consistent throughout the paper. Panel (**A**) shows the structures of the F-ATPase from *Escherichia coli* (PDB ID 6OQW [22]), the A-ATPase from *Thermus thermophilus* (PDB ID 6QUM [4]), and the V-ATPase from yeast (PDB 7TMS [23]. Regions of proteins that can be attributed to certain protein family clusters (Clusters of Orthologous Groups, COGs) are colored. Eukaryotic proteins that do not correspond to any prokaryotic homolog are shown in gray. Panel (**B**) shows typical operons of F- and A-type ATP synthases from *E. coli* and *T. thermophilus,* as well as the COGs which correspond to each protein. Color scheme is the same as in panel **(A)**. Panel **(C)** shows the structures of Na⁺-binding ***c***-rings of bacterial F- and A-type ATPases: the ***c****-*rings from *Ilyobacter tartaricus* (PDB ID 1YCE [19]) and *Fusobacterium nucleatum* (PDB ID 3ZK1 [24]) belong to F-ATPases, whereas *Enterococcus hirae* has an A-type ATPase (PDB ID 2BL2 [20]).

Rotary ATPases are built of two distinct parts: a membrane anchored part (F_O_, A_O_ and V_O_, respectively) and a cytoplasmic part (F_1_, A_1_ and V_1_, respectively), see Fig. 1A, B.

The core of the cytoplasmic part is a doughnut-shaped hexamer made up of alternating catalytic (β-subunits in bacteria and A-subunits in archaea) and non-catalytic (α-subunits in bacteria and B-subunits in archaea) subunits, see Fig. 1A. These subunits are homologous and contain so-called <u>P</u>-loops (Walker A motifs) that can bind the **<u>p</u>**hosphate groups of ADP or ATP [15–17]. Therefore, we will refer to both the catalytic and non-catalytic subunits together as “P-loop subunits” from now on. The central rotating shaft, made of an elongated coiled-coil (γ-subunit in bacteria and D-subunit in archaea), sequentially interacts with the catalytic subunits and promotes either the ATP synthesis or its hydrolysis, depending on the direction of rotation, see Fig. 1A. Hence, a full rotation of the central shaft is accompanied by either hydrolysis or synthesis of three ATP molecules by the catalytic subunits [7,8,10].

The membrane part includes a membrane-embedded subunit ***a*** (hereafter, unless indicated otherwise, we use the same membrane subunit names as in F_O_F_1_-ATPase of *E. coli* also for A_O_A_1_- and V_O_V_1_-ATPases). The ***a***-subunit interacts with a rotating ring of highly hydrophobic ‘proteolipid’ ***c***-subunits, see Fig. 1A. In the F-type ATPases, these ***c***-rings are predominantly built of hairpin-like subunits, each of which is formed by two transmembrane helices. In archaea and eukaryotes, the cognate ***c***-subunits may have more helices owing to gene duplications; the typical numbers are two and four, see Fig. 1A, C and [4,8,18–21]. The ions are translocated along the interface between the rotating ***c***-ring and the ***a***-subunit.

The peripheral stalks, each built of two elongated subunits, connect the P-loop hexamer with the membrane part, see Fig. 1A. Consequently, the catalytic heterohexamer, the outer stalk(s), and the ***a***-subunit make the ‘stator’, while the inner shaft and the ***c***-oligomer form the “rotor”.

While three ATP or ADP molecules are processed per one full turnover, the number of ions translocated depends on the number of ion-binding sites of the ***c***-ring and varies greatly due to the elastic coupling between the membrane and the cytoplasmic parts [7,25,26]. The variability of the proteolipid ring is constrained by the trade-off between the magnitude of the Δμ̅ H^+^ or Δμ̅ Na^+^ (usually about 200 meV ∼ 20 kJ/mol in prokaryotes) and the free energy of ATP synthesis in the cell (about 50-60 kJ/mol ∼ 500-600 meV) [12,13]. The smallest number of ***c***-subunits known is eight; it is found in animal mitochondria [27], which appear to produce ATP most efficiently.

These structural traits are common to all rotary membrane ATPases, which may differ in other ways, as discussed below.

In F-ATPases, the two ***b-***subunits of the outer stalk are anchored in the membrane by their N-terminal hydrophobic helices interacting with those of the ***a***-subunit (Fig. 1A). The two outer stalks of A_O_A_1_-ATPases and the three outer stalks of V_O_V_1_-ATPases, each consisting of E- and G-subunits, lack such hydrophobic membrane-anchored helices. Instead, they attach to the cytoplasmic protrusions of the ***a-***subunit and also to the subunit C in V-ATPases, see Fig. 1A.

Most F-ATPases have a δ-subunit on the top that fastens the interaction between the *b*-subunits and the α_3_β_3_ hexamer, see Fig.1A. The A- and V-ATPases lack an additional subunit in this position.

Also, F-ATPases contain the ε-subunit that rotates with the γ-subunit and ***c***-ring, see Fig. 1A [28]. The central stalks of A- and V-ATPases can contain a small F-subunit in a similar position, see Fig. 1A [4,5,29–31].

As seen in Fig. 1A, the A- and V-ATPases have a conical ***d***-subunit that sits atop the ***c***-ring and serves as a socket for the central shaft [4,5,29–31]. This subunit has no functional counterpart in F-ATPases.

Eukaryotic V-ATPases, as well as F-ATPases of mitochondria, have further additional subunits absent in their prokaryotic counterparts, see [7,9] for reviews.

The overall bizarreness of rotary ATPases has long raised questions about their evolutionary origin. Specifically, Walker has suggested that the membrane and cytoplasmic parts may have evolved separately initially, as a hexameric ATPase and a membrane transport pore, respectively. Only later, after recruiting the outer stalk(s), they became the rotary machine capable of ATP synthesis [32,33].

Cross and Taiz have hypothesized that the original membrane complex served as a proton-translocating ATPase with six identical catalytic subunits. They proposed that the loss of half of the catalytic sites was necessary to make the ATP synthesis by the ion gradient thermodynamically feasible [34]. This scenario got support when it was found that the P-loop subunits of membrane rotary ATPases are homologous to the *FliI* subunits of the homohexameric basal ATPase of bacterial flagella. In addition, similarities were found between the flagellar *FliH* subunit and the outer stalk subunits of both F- and A/V-ATPases [35]. Together, *FliI* and *FliH* are involved in transport and partially unfolding of flagellin before its export through the hollow bacterial flagellum [36].

Building on this evidence, we previously hypothesized that ion-transporting rotary membrane ATPases evolved from a protein export pump, in which the transported protein became lodged in the pore-forming membrane complex and caused it to rotate. Recalling Cross and Taiz’s ideas [34], we hypothesized that this ancestral ATPase had a homohexameric catalytic part. For reasons of symmetry, we suggested that the hexamer was supported by two outer stalks [37].

Crystal structures of Na^+^-translocating ***c***-rings, as resolved for the F-ATPase [19] and A-ATPase [20], revealed Na^+^ ions bound by sets of specific residues (Fig. 1C). These residues were found to be homologous in F- and A-type ATPases [38,39]. Because Na^+^-translocating ATPases with similar Na^+^-binding residue sets were found in many bacterial and archaeal branches of the phylogenetic tree, it was inferred that the ancestral membrane rotary ATPase of LUCA was a Na^+^-binding ATPase [38,39].

The ancestral rotary ATPase likely served as a Na^+^-export pump [39]. Earliest cells required such pumps because the protein synthesis system — arguably the oldest cell machinery [40–42] — demands approximately ten times more K⁺ ions than Na⁺ ions, which contrasts with the prevalence of sodium in both fresh and seawater [43–45].

Remarkably, mutating Na^+^-binding residues in rotary ATPases results in loss of Na⁺-binding ability, though proton-transport ability is usually preserved [2]. In many organisms, e.g. in *E. coli*, proton translocation by ***c***-subunits requires just one carboxylic acid residue in the middle of their second transmembrane helix (Fig. 1A). Consequently, it has been suggested that the transition to H⁺-specific rotary ATPases occurred independently — through the sporadic loss of Na⁺-binding residues — in distinct bacteria and archaea after their membranes became proton-tight [38,39]. This suggestion aligns with the observation that different Na⁺-binding residues were lost in different bacterial lineages [46]. H⁺-dependent energetics may have became particularly advantageous after the oxygenation of the atmosphere approximately 2.5 billion years ago [46,47].

The evolutionarily primacy of Na^+^-translocating rotary ATPases is supported by the identification of a subfamily of primitive F-ATPases with mostly Na^+^-translocating ***c***-rings [48]. These so-called N-ATPases lack the δ subunit; instead, the ***b*** subunit is elongated by a segment similar to the C-terminal part of the δ subunit in other F-ATPases. Remarkably, the ***b***-subunit of N-ATPases shows similarity with the cognate subunit E of A-ATPases and the flagellar *fliH*-subunit [35,49]. It has been suggested that N-ATPases, almost always found alongside the “main” F-or A-ATPases of prokaryotic cells, may have a specialized function, e.g. of Na^+^-exporting pumps [48]. Takabe and his colleagues showed that expressing the N-ATPase operon from *Acaryochloris marina* in the heterologous cyanobacterium *Synechococcus* sp. PCC 7942 increased the salt stress tolerance. The same authors demonstrated the ability of this N-ATPase for Na^+^-coupled synthesis and hydrolysis of ATP when incorporated into membrane vesicles [50].

Further progress in understanding the evolution of rotary ATPases was driven primarily by advances in obtaining their structures from various organisms, as covered in several comprehensive reviews [4,5,7–11,29–31,51].

Yet, over the recent few years, exciting new functional, structural and genomic data have challenged the established views on rotary membrane ATPases.

Specifically, Matzke and colleagues found that the similarity between ion-translocating rotary ATPases and the basal ATPases of bacterial flagella involves at least three proteins, for which not only the amino acid sequences but also the gene order is conserved in some cases [49].

Two recent articles updated the distribution of rotary membrane ATPases in light of the surge in sequenced genomes. The analysis revealed numerous instances of lateral transfer of rotary ATPase genes between and within the bacterial and archaeal domains [52,53]. The apparent asymmetry of this transfer — with A-ATPase transfers to bacteria being much more frequent than only few — if not only one — F/N-ATPase transfer to archaea — led to the consideration of several alternative evolutionary scenarios. One such scenario is that both types of ATPases were present in LUCA, followed by the almost complete loss of F-ATPases in the archaeal lineage [52].

Noji and his colleagues created more space at the top of the α_3_β_3_ hexamer of the F-ATPase of *Bacillus* PS3 by truncating the N-terminus of the δ-subunit and fusing its C-terminus to the α-subunit. This allowed up to three peripheral stalks (***ab*δ_truncated_** complexes) to bind to non-catalytic α-subunits. Each stalk interacts with the *c*-ring via its own ion-translocating ***a***-subunit. The authors argue that this configuration can triple the number of protons translocated per one full rotation, enabling ATP synthesis at low membrane potential [54].

Meanwhile, Xu and her colleagues obtained the cryo-EM structure of the F-ATPase from the early-branching bacterium *Chloroflexus aurantiacus*. They discovered that this F-ATPase has two outer stalks, each of which interacts with the ***c***_10_-ring via a separate ***a***-subunit. In this bacterium, the synthesis of three ATP molecules is apparently driven by the translocation of 20 protons. The authors argue that the smaller δ-subunit of *Chl. aurantiacus* leaves enough space atop the α_3_β3 hexamer for the binding of an additional outer stalk [55].

At the same time, Osipov and his colleagues presented the results of computational approach based on AlphaFold that enabled predicting the stoichiometry of all homo-oligomeric *c*-rings of F-type ATPases based on their amino acid sequences. After validating their approach on the available experimental data, the authors estimated the number of ***c***-subunits in over one thousand representative genomes and predicted the existence of native ***c***-rings built of up to 27 subunits. The authors hypothesized that such rings might be useful at low transmembrane potentials [56].

These seminal findings from recent years are not easily reconciled with earlier evolutionary scenarios, which prompted us to revisit the evolution of rotary ATPases.

Here, we combined phylogenomic tools with comparative structural analysis, which allowed us to adjust our earlier evolutionary scenarios [37–39] to align with the new data. We posit that the evolution of rotary ATPases was driven by the need to increase the net ATP production, albeit at the expense of mechanistic stability in the case of F-ATPases.

## 2. Methods

### 2.1. Sequence samples

We used a complete representative sample of 2296 prokaryotic genomes covered by the **<u>C</u>**lusters of **<u>O</u>**rthologous **<u>G</u>**roups of proteins (COG) database [57]. By confining the sample to these previously selected high-quality genomes, we eschew reliance on incomplete and inadequately annotated genomes. Yet, it must be noted that confining the number of genomes entails the risk of overlooking rare variants of rotary ATPases. For β-subunits (COG0055), δ-subunits (COG0712) and ***c***-subunits (COG0636) of F-type ATPases we sampled both sequences originally attributed to the respective COGs [57] and those identified with our set of profile HMMs for COGs [58]. We grouped β, δ and ***c***-subunits based on their genomic neighborhoods, identified orphaned β and δ subunits, mostly due to a missing gene in the genome annotation, and corrected these cases. Sequences of A-subunits (COG1155) and flagellar ATPase FliI subunits (COG1157) were sampled from the COG database.

### 2.2. Mapping predictions for the number of membrane c-subunits on the data on β- and δ-subunits

In their work, Osipov and his colleagues took ***c***-subunit sequences from 58,732 bacterial species and combined them into 1,571 clusters. One sequence was selected from each cluster for further evaluation and prediction of the number of ***c***-subunits in the ring. [56]. We managed to obtain 53,808 out of these sequences from the UniProt, compared them with the set of ***c***-subunits attributed to the sampled β-subunits using the BLAST+ program [59], and identified 1,634 proteins that were 100% identical. The numbers of ***c***-subunits, as predicted by Osipov and colleagues for these proteins, were mapped to our data.

### 2.3. Prediction of ion specificity of c-subunits from their sequences

We inspected sequences of the ***c***-subunits and predicted their ion specificity based on the presence/absence of a sodium-binding motif [38]. Specifically, the enzyme was considered Na^+^-translocating if its ***c****-*subunit met the following two conditions:

1. A Glu residue (E^2^) is present in the middle of the second helix of the single ***c***-subunit hairpin (for double-hairpin ***c***-subunits this is required only for the second hairpin), while the first helix contains Glu or Gln in the middle (E^1^ or Q^1^);
2. The second helix of a ***c***-subunit hairpin contains a pair of Na^+^-binding neutral residues (Ser, Thr or Gln) after E^2^ (E^2^+1 and E^2^+2 positions), and a Tyr residue is in the E^2^+5 position (for double-hairpin ***c***-subunits this is required only for the first hairpin). Thus, the motif required for the second helix of a single-hairpin ***c***-subunits is E[STQ][STQ]xxY.

When a single S/T/Q residue was missing in the E^2^+2 position, but all other possible Na^+^ ligands were present, we considered a possibility of Na^+^ translocation, as demonstrated for the F-type ATP synthase of *Synechocystis* sp. PCC 6803 at pH > 8.0 [60].

Some double-hairpin proteins contained potential Na^+^-binding residues in positions that were uncommon compared to those in structurally studied systems. For example, an [STQ][STQ]xxY motif could be present in the second hairpin, which also occurs in several multi-hairpin structures. We marked such motifs as "H^+^/Na^+^" to distinguish them from proteins without any possible Na⁺-binding site.

### 2.4. Obtaining additional sets of sequences

We complemented the above data sets by sequences of the β-, δ- and ***c***-subunits of several F-type ATPases absent in the COG database. For these F-ATPases, we used the number of ***c***-subunits, as experimentally determined from structural data, as reference values. For that, we analyzed the genomes of *Bacillus sp.* PS3 (GCF_005154805.1), *Clostridium paradoxum* (GCF_001584565.1), *Arthrospira platensis* HN01 (GCF_026000695.1), *Mycolicibacterium phlei* (GCF_900453675.1), *Chloroflexus aurantiacus* J-10-fl (GCF_000018865.1) and *Burkholderia pseudomallei* S13 (GCF_000152365.1). For *Propionigenium modestum* and *Ilyobacter tartaricus,* we used the respective operon sequences (GenBank records X58461 and AF522463 respectively), since their complete genome assemblies are still not available.

To examine unusual genomic neighborhoods of β-subunit homologs from the Candidate Phyla Radiation (CPR) bacteria, we obtained from this phylum additional 12264 genome assemblies with predicted genes and assigned them to COGs using our set of profile hidden Markov models (HMMs) for COGs [58]. Since the resulting sample of β-subunits of F-type ATPases (COG0055), A-subunits of A-type ATPases (COG1155), and flagellar ATPases FliI (COG1157) was large, we reduced it using the UCLAST software [61] at 90% identity threshold.

### 2.5. Assessment of the thermophilicity of A-ATPase hosts

Data on the thermophily of the species with A-type ATPases were extracted from the IMG and BacDive databases [62,63] or were checked manually through the literature.

### 2.6. Multiple alignment and phylogenetic trees construction

Sequences were aligned using Muscle 3.8.31 [64]. Phylogenetic trees were obtained using IQ-TREE version 3 [65] with the ultrafast bootstrap support [66] and FastME version 2 [67] followed by the transfer bootstrap expectation calculation [68] by the BOOSTER (https://booster.pasteur.fr/). Further parameters of tree construction are provided in the figure captions. Attribution of species to their taxonomic groups was done using the NCBI Taxonomy database [69]. Genomic neighborhoods were visualized using the home-made pyCOGNAT software (https://github.com/udavdasha/pyCOGNAT).

## 3. Results

### 3.1. Phylogenomic analysis of F-ATPases

Figure 2 shows the phylogenetic tree for catalytic β-subunits of F-type ATPases superimposed over the data from the same organisms regarding (i) the number of subunits in their ***c***-rings, as established by structural methods and/or predicted by Osipov and colleagues [56], as well as (ii) the length of their δ-subunits. The “rectangular” version of the same tree, including the genomic neighborhoods of all amalyzed β-subunits, is given in Supplementary File 1. Note that this tree does not represent the full diversity of F-type ATPases; it contains only those for which Osipov and colleagues predicted the number of subunits in the ***c***-rings [56] and is further restricted to ATPases from representative, well-assembled genomes included in the COG database [57].

**Figure 2.**
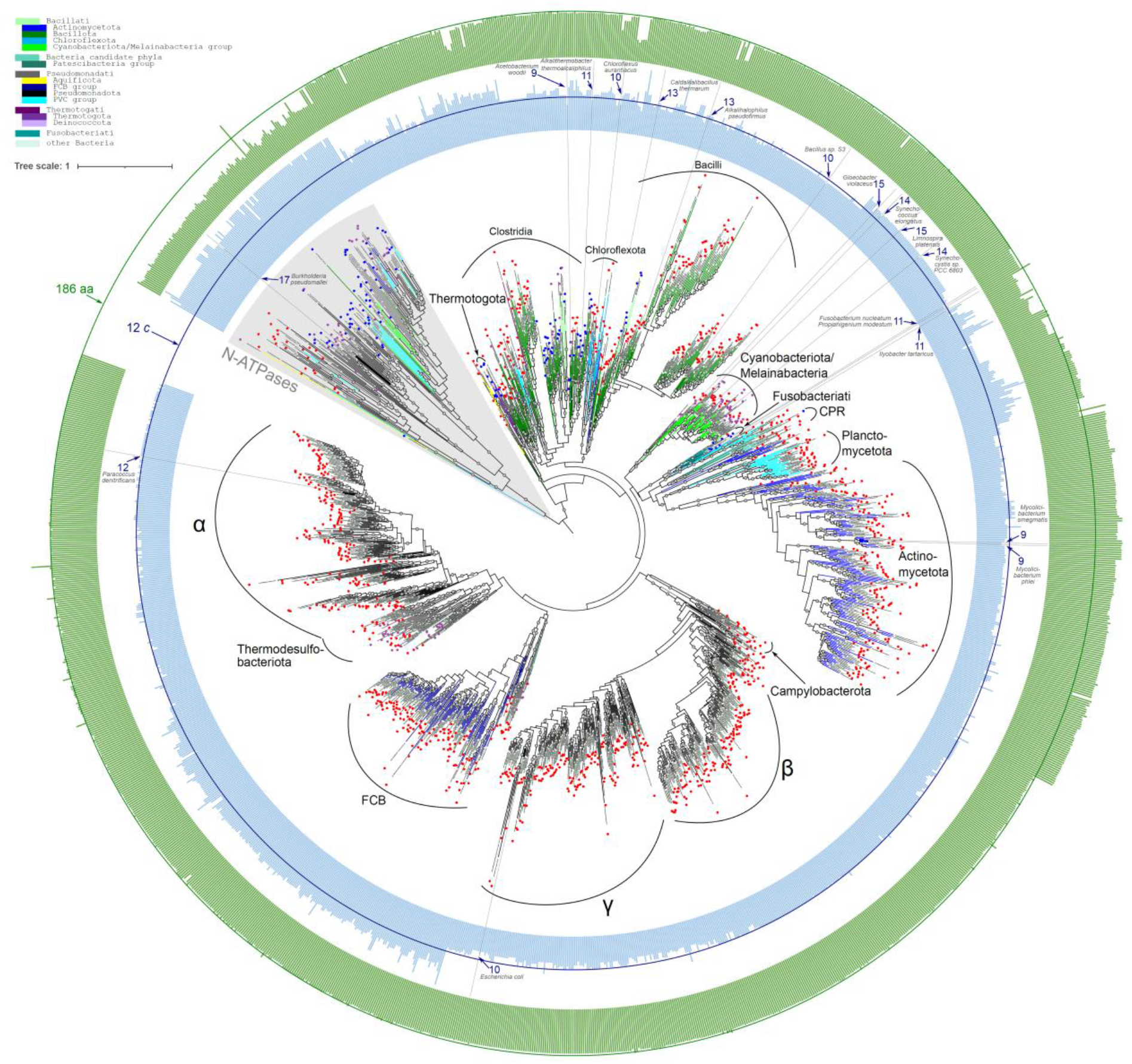
Phylogenetic tree for the β-subunits of F-type ATPases. The tree is superimposed over the data from same organisms regarding (i) the number of subunits in their ***c***-rings, as established by structural methods and/or predicted by Osipov and colleagues [56], as well as (ii) the length of their δ-subunits. [56]. The multiple alignment contained 1484 sequences and 893 positions. The tree was constructed with the IQ-TREE, ModelFinder [73] yielding the LG+I+R10 evolutionary model for this dataset. Ultrafast bootstrap approximation above 95 is shown with gray circles on the branches. The branches are colored according to the taxonomy of the respective species. The color code for taxonomy is shown in the top left corner. The internal blue ring shows the predicted number of ***c***-ring subunits according to [56], whereas the experimentally established numbers of ***c***-ring subunits are directly marked by gray lines and indicated by arrows. The external green ring shows the lengths of the δ-subunits (or, in cases of fusion between the δ-subunit and either the α-or ***b***-subunit, the length of the fused protein segment not covered by the respective α-or ***b***-subunit HMM profile). The prediction of the ion specificity of the ***c***-subunits is shown by circles at the ends of branches. Red circles show H⁺ specificity, blue circles show Na⁺ specificity, and purple circles indicate nearly complete Na⁺-binding sites with one missing residue.

In light of the data by Noji et al. considered in the Introduction [54], we were interested in the relationship between the length of the δ-subunit and the predicted number of subunits in the ***c***-ring. Generally, ***c***-rings with many (∼20) subunits are associated with shorter δ-subunits — or protein segments homologous to them in the organisms lacking separate δ-subunits. For example, N-type ATPases have no separate δ-subunits; however, the C-terminus of their ***b***-subunit contains a segment homologous to the С-terminal part of a separate δ-subunit. This segment is much shorter than a typical δ-subunit, and its divergence often hinders its attribution to the respective COG0712 (Fig. 1B), even with a non-strict e-value of 0.01 (see Supplementary Figure 1). The only experimentally established value for the number of ***c***-subunits in an N-type ATPase is 17 for *Burkholderia pseudomallei* S13[70]. Yet, the numbers of ***c***-subunits predicted for N-ATPases are mostly over 20. Our data reveal fusions between subunits ***b*** and δ, as well as between the δ-subunit and the non-catalytic α-subunit downstream in the operon (where, as expected, a region homologous to the δ-subunit or only to its C-terminal part is located at the N-terminus of the α-subunit, see Fig. 1B and Supplementary Figure 1). These fusions are accompanied by the shortening of the segment homologous to the δ-subunit. Another atypical case is the ATP synthase of Actinomycetota, where the δ subunit is much longer and the predicted number of ***c*** subunits is slightly lower than in other complexes. Experimentally studied F-ATPases from *Mycolicibacterium smegmatis* MC2 155 and *M. phlei* contain only nine subunits in their ***c***-rings [71,72].

### 3.2. Phylogenomic analysis of A-ATPases

To clarify why there are many more bacteria with A-ATPases than archaea with F-ATPases [52,53], we constructed the phylogenetic tree for catalytic A-subunit of A-type ATPases. The tree is shown in Fig. 3; its complete version with genomic neighborhoods of the catalytic A-subunits is given in Supplementary Figure S2.

**Figure 3.**
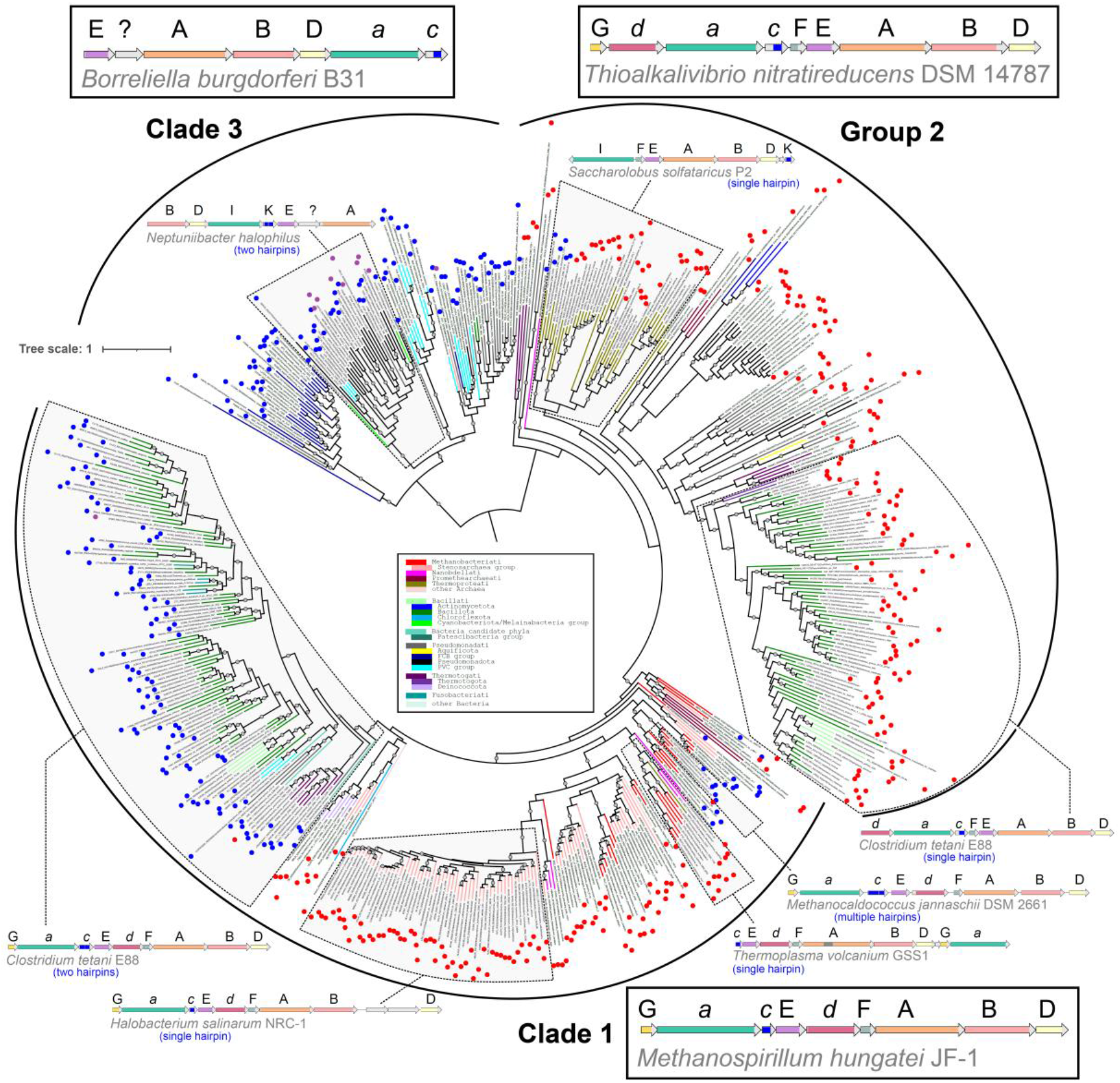
Phylogenetic tree for catalytic A-subunits of rotary membrane A-type ATPases (COG1155). The complete version with all genomic neighborhoods of the catalytic A-subunit is given in Supplementary Figure S2. The multiple alignment contained 548 sequences and 1650 positions (including regions of intein insertions which occur in some proteins). The tree was constructed with the IQ-TREE, ModelFinder [73], which yielded the LG+I+R10 evolutionary model for this dataset. Ultrafast bootstrap approximation above 95 is shown with gray circles on the branches. The ultrafast bootstrap approximation above 95% (from 1000 calculations) is shown by gray circles on the branches. The branches are colored according to the taxonomy of the respective species. The color code for taxonomy is shown in the top left corner of Fig. 2. The genomic neighborhoods typical for the boxed segments of clades 1, 3 and group 2 are framed. The prediction of the ion specificity for the ***c***-subunits is shown by circles at the ends of branches. Red circles indicate H⁺ specificity, blue circles show Na⁺ specificity, and purple circles indicate nearly complete Na⁺-binding sites with one missing residue.

The tree clearly splits into three segments, which differ in both the sequence of the catalytic subunit and the gene operon structure (see Fig. 3 for typical operon structures and Supplementary Figure S2 for the complete phylogenetic tree with structures of all operons).

Clade 1 contains most archaeal sequences, including representatives of the phyla Thermoplasmatota, Halobacteriota, and Methanobacteriota, as well as enzymes from bacterial phyla such as Firmicutes and Deinococci. These enzyme complexes all exhibit operon structures typical of archaeal A-ATPases. In these operon structures, the P-loop subunit genes are separated from the genes of the outer stalk-forming E subunit by the genes of subunits ***d*** and F, which together with the D subunit form the central stalk.

Group of clades 2 also contains enzymes from both archaea and bacteria. The bacterial representatives include Firmicutes, Synergistota, Spirochaetota, and Proteobacteria. Archaea are represented by only two kingdoms: Promethearchaeota and Thermoproteota. However, the enzyme complexes of group 2 have a different operon structure. Here, the gene of the outer stalk-forming E subunit adjoins the genes of the P-loop subunits. The genes of the ***d*** and F subunits are located elsewhere in the operon or are not identifiable in many cases.

Clade 3 consists mostly of bacterial sequences, except for *Methanolinea* sp. The host bacteria belong predominantly to Spirochaetota and Proteobacteria, which are also present in clade 1 and group of clades 2. Chlamydiae and Bacteroidetes are also present. Almost all A-ATPases in this clade are Na⁺-dependent (see Fig. 3). Furthermore, clade 3 is subdivided into two branches. These branches differ from one another in that the second half of the operon in one branch has shifted to the beginning in the second branch. They also differ in length and, consequently, in the number of hydrophobic helices in the membrane ***c***-subunit, see Supplementary Fig. S2.

The gene operons of clade 3 and some branches of group 2 do not contain the F subunit gene, see Supplementary Fig. S2. Another ambiguity is coupled with the ***d***-subunit. Its gene is absent from some group 2 A-ATPase operons. Additionally, the gene at the respective position is recognized as a ***d***-subunit in only a few clade 3 organisms, see Supplementary Fig. S2. This protein is approximately 50–100 amino acid residues shorter than a typical ***d***-subunit. Its alignment with "canonical" ***d***-subunits suggests that they are different proteins.

Archaea are generally more heat-tolerant than bacteria. This implies that their proteins are more thermostable [21]. Consequently, particular bacteria may adapt to higher temperatures by acquiring A-ATPases. If so, thermophilic bacteria may prevail among bacteria with A-ATPases. Therefore, we checked whether the presence of A-ATPases in some bacteria is due to the higher thermostability of A-ATPases compared to F-ATPases. Indeed, many bacteria possessing A-ATPases are thermophiles.

### 3.3. Phylogenomic analysis of the catalytic subunits of membrane rotary ATPases and flagellar ATPase

Figure 4 shows the phylogenetic tree constructed for the sequences of catalytic subunits of prokaryotic rotary ATPases and their closest homologs, namely the catalytic subunits FliI of the bacterial flagellar ATPase. The complete version of the tree with genomic neighborhoods of all examined catalytic subunits is given in Supplementary Figure S3. The branch lengths indicate that the catalytic subunits of archaeal and bacterial ATP synthases are slightly more dissimilar from each other than from the flagellar FliI ATPase. This finding is consistent with our previous observations [74]. We discuss the probable cause of this strong divergence in the "Discussion" section.

**Figure 4.**
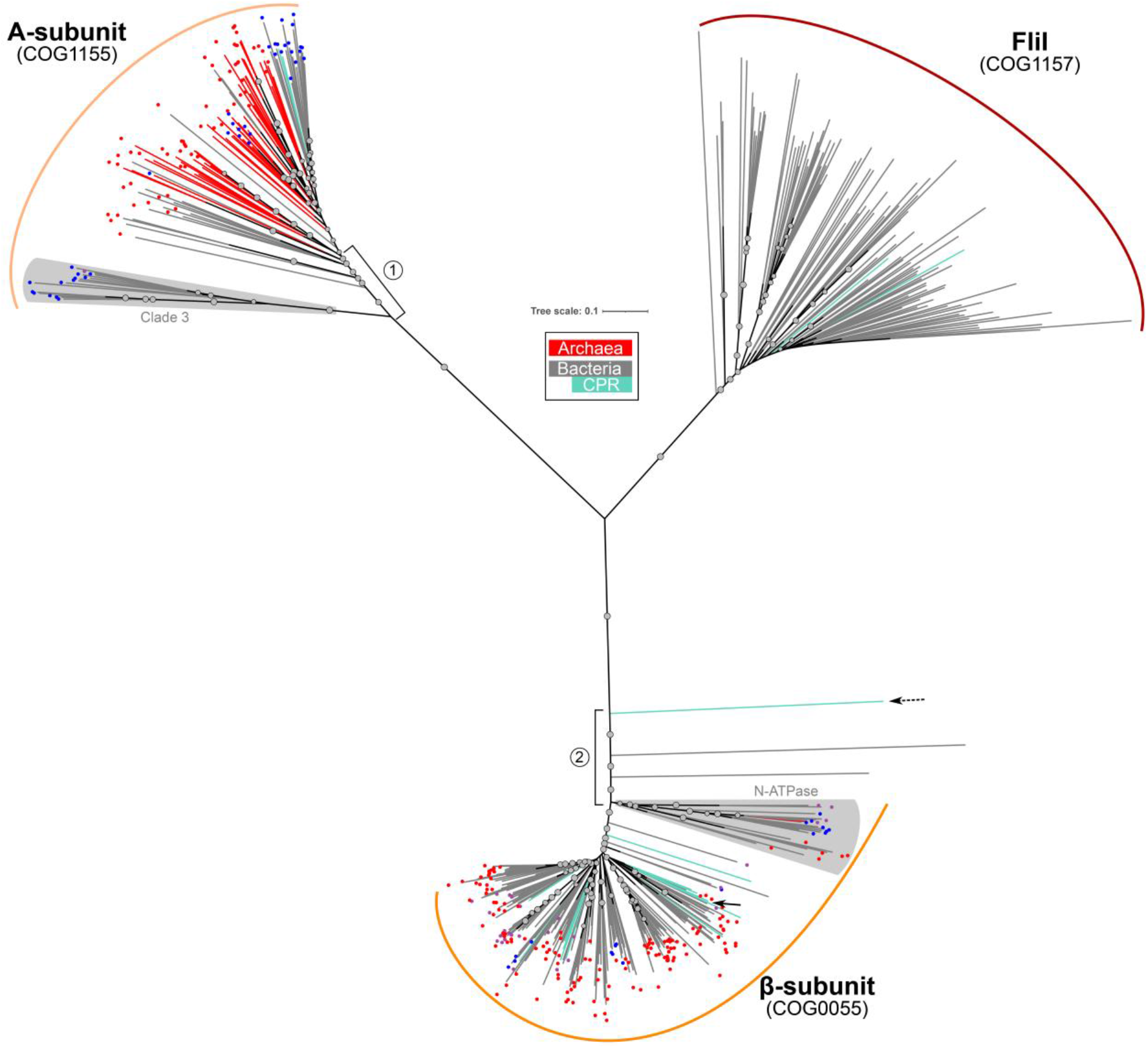
Phylogenetic tree of the catalytic β-subunits of bacterial F-ATPases (COG0055), the catalytic A-subunits of archaeal A-ATPases (COG1155), and the homologous flagellar ATPase FliI (COG1157). The complete version of the three with all genomic neighborhoods of the catalytic subunits is given in Supplementary Figure S3. The tree was constructed using a reduced representative sample of 275 complete genomes included in the latest COG database version [57]. We used FastME with the LG evolutionary model; TBE support values above 70 for 100 replicates are shown as circles. The tree was visualized in iTOL (https://itol.embl.de/). The early-branching A-subunits of A-ATPases are labeled (1), and the early-branching β-subunits of F-ATPases are labeled (2). Clades with unusual genomic contexts, such as N-ATPases and an early-diverging clade 3 within A-ATPases, are highlighted with a light gray background. The ionic specificity of A- and F-ATPases is labeled as in Fig. 2 and 3.

As shown in Fig. 4, the main clades of catalytic subunits of A- and F-ATPases are preceded by early branching groups on the phylogenetic tree. For F-ATPases, the largest early-branching clade is that of the N-ATPases, most of which are Na⁺-translocating [48]. In the case of A-ATPases, the Clade 3 of mostly Na⁺-translocating A-ATPases branches first, see the previous section.

### 3.4. Comparison of operon structures for A-ATPases, F-ATPases, and basal ATPases of bacterial flagella

Figure 5 compares the gene operon structures of flagellar ATPases with those of A- and F-ATPases, focusing on their basal clades, shown in Fig. 4.

**Figure 5.**
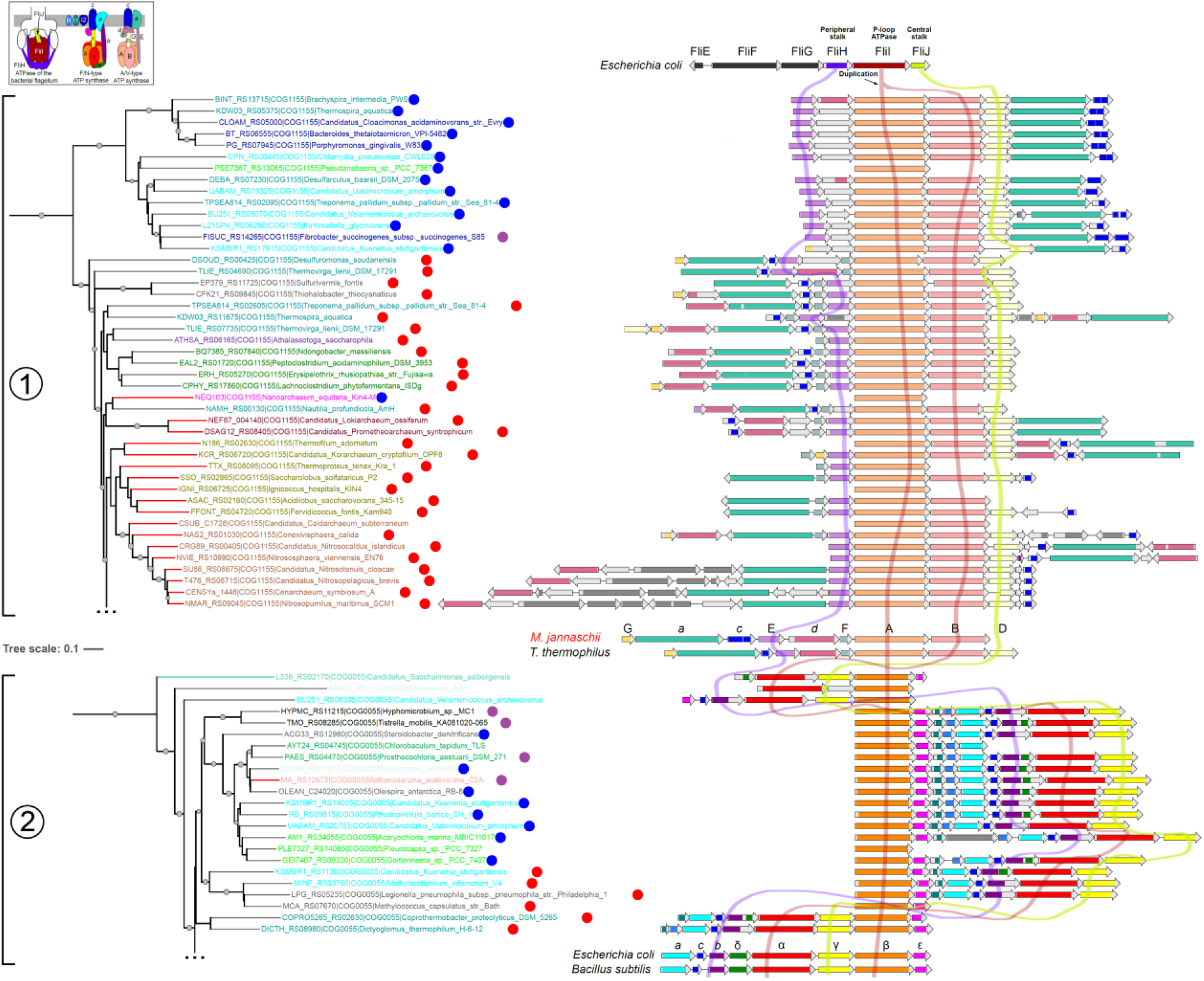
Genomic contexts of bacterial flagellar ATPase and the catalytic subunits of rotary membrane ATPases. The operon structures of early branching, "basal" A- and F-ATPases (see the text, Fig. 4 and Supplementary Figure S3) are compared to those of the flagellar ATPase and the typical genomic contexts of A-ATPases (from the archaeon *Methanocaldococcus jannaschii* and the bacterium *Thermus thermophilus*) and F-ATPases (from *Escherichia coli* and *Bacillus subtilis*).

Previously, Matzke et al. observed that the gene order of the three flagellar ATPase genes (fliHIJ) is similar to the arrangement of their homologs in certain membrane ATP synthases. These authors considered this order to be evolutionarily primary [49]. Our data show that rotary ATPases with a “flagellar” gene order belong to the early-branching “basal” clades highlighted in Fig. 4. In many of these ATPases, the genes are arranged in this order: outer stalk – P-loop subunit(s) – central stalk. In contrast, the majority of A- and F-ATPases with a more canonical gene order have the outer stalk subunit (E-or ***b***-subunit, respectively) separated from the P-loop subunit(s) by additional subunits (***d*** and F in A-ATPases and δ in F-ATPases). This results in a deviation from the characteristic gene order of bacterial flagellar ATPases.

## 4. Discussion

### 4.1. Relation between the length of the δ-subunit and the (predicted) number of subunits in the c-rings of F-ATPases

As shown in Fig. 2 and Supplementary Figure S1, the predicted large number of ***c***-subunits (>20) parallels the short length of the δ-subunit or its homologous domain within the ***b***-subunits of δ-subunit-lacking bacteria (< 130 amino acids). This pattern is evident among all N-ATPases and several groups of F-ATPases with short δ-subunits belonging to CPR bacteria, Verrumicrobiota, Chloroflexota, and Bacilliota. Furthermore, F-ATPases with short δ-subunits and a "normal" number of c-subunits (≤15) are also seen, as in the previously mentioned *Chl. Aurantiacus* [55]. Interestingly, short δ-subunits are primarily observed in early-branching bacterial phyla.

As discussed in the Introduction, the presence of a short δ subunit may allow the binding of additional outer stalks [54,55]. Therefore, the data in Fig. 2 and Supplementary Figure S1 can be interpreted as evidence that the N-ATPases, which lack the δ-subunit, have more than one outer stalk in combination with ∼20 ***c***-subunits. Under assumption that all their outer stalks are ion-translocating as in *Chl. aurantiacus* [55], the synthesis/hydrolysis of three ATP molecules upon a full rotation of an N-ATPases would be coupled with translocation of ∼40 ions if they have two outer stalks (as in *Chl. aurantiacus* [55]) or of ∼ 60 ions if they have three outer stalks as in genetically modified *Bacillus* PS3 [54].

Then, however, the optimal operating mode of the N-ATPases /synthases would correspond to values of ΔµH⁺ or ΔµNa⁺ of ∼ 50 mV. This may explain why N-ATPases function as additional ATPases. These complexes could be useful and efficient as ATPases or ATP synthases at low membrane potentials. For example, this could occur in the absence of light (in phototrophs) or food. The common occurrence of N-ATPases among phototrophic bacteria [48] indicates their involvement during dark periods when the membrane potential may drop. Therefore, N-ATPases likely serve as standby ATP synthases for cases of membrane potential drop in their present-day hosts.

Consequently, F-ATPases with short δ-subunits may also have more than one outer stalk, as seen in *Chl. aurantiacus* [55]. F-ATPases with more than one ion-translocating stalk may be useful for organisms with leaky membranes that are poor at maintaining high membrane potentials. This may be the case in some thermophilic bacteria and some CPR bacteria with many lysolipids in their membranes [75].

Named reasons may explain the presence of the second stalk in the F-ATPase of *C. aurantiacus* [55], which is both phototrophic and thermophilic, living in hot springs [76].

It is tempting to speculate that F-ATPases evolved from complexes with more than one outer stalk and many ***c***-subunits, through complexes with two outer stalks and fewer ***c***-subunits, and finally to the ‘canonical’ F-ATPase with one outer stalk and 15 or fewer ***c***-subunits. This point will be addressed in more detail below.

### 4.2. Was the lateral transfer of A-ATPases to bacteria driven by their high thermal stability?

Our phylogenomic analysis of A-ATPases revealed three distinct groups, all containing archaea and bacteria. In two of these groups, distinct archaeal and bacterial branches could be identified (Fig. 3, Supplementary Fig. S2). These results suggest that A-ATPases spread among bacteria at least twice, likely following two or more lateral gene transfer events. Our analysis revealed that many bacteria using A-ATPases appear to be thermophilic or hyperthermophilic. Thus, the abundance of A-ATPases in bacteria can be explained by the greater thermal stability of archaeal A-ATPases [21]. This trait makes A-ATPases an asset for thermophilic bacteria, particularly those living under energy-limiting conditions. Since our data indicate at least two "waves" of lateral transfer of A-ATPases to bacteria, it is unlikely that LUCA contained both F-ATPases and A-ATPases, which was discussed in [52].

### 4.3. Comparative phylogenomic analysis of the catalytic subunits of the rotary membrane ATPases and the basal ATPase of bacterial flagella

Fig. 4 shows that the catalytic subunits of F- and A-ATP synthases differ more from each other than from flagellar ATPases. This divergence may be related to the loss of half of the catalytic sites in the ATP synthases. Indeed, P-loop ATPases often form ring-like oligomers wherein the catalytic reaction within the subunit with an already bound substrate molecule is promoted by the binding of another substrate molecule to the adjacent subunit [77–79]. Closing the catalytic site in the adjacent subunit triggers a conformational change that results in the insertion of a stimulating moiety (usually an arginine “finger”) into the catalytic site of the subunit with an already-bound ATP. Thus, the energy of substrate binding in one site is used to promote catalysis in the other site [79,80]. Such cooperativity is found in flagellar homohexameric ATPases [81] and was likely inherent to the homohexameric ancestor of A- and F-ATPases. However, the transition from a homohexameric ancestral rotary ATPase to the heterohexameric A- and F-ATPases, in which the cooperative interaction between the catalytic subunits is transmitted via an intervening, bulky, noncatalytic subunit, must have required significant structural changes.

Furthermore, the significant divergence of orthologous catalytic subunits in Fig. 4 suggests that the separation of catalytic and non-catalytic subunits may have happened independently in F/N- and A/V-ATPases. This may result either from two distinct gene duplication events or from two independent losses of the catalytic function after a single gene duplication. Such cases are known (see e.g. [82,83]). Even in F-ATPases, the ***b***-subunit-coding gene has undergone several independent duplications, resulting in distinct functional divergences between the respective ***b*** and ***b’*** subunits [84].

This suggestion gets support from a comparative structure analysis, see Fig. 6. The P-loop subunits of membrane rotary ATPases belong to the RecA/F_1_ class of P-loop nucleotide triphosphatases (NTPases) [17]. As argued elsewhere, all P-loop NTPases have the same catalytic site arrangement [15–17,85–87]. Its obligatory elements are the Walker A (WA) and Walker B (WB) motifs [15]. The WA motif (GxxxxGK[S/T]) (i) forms the P-loop that „embraces” the tri**<u>p</u>**hosphate moiety, (ii) compensates for its negative charge through a strictly conserved Lys residue (Lys^WA^) and several backbone amino groups, and (iii) provides a strictly conserved Ser/Thr residue (Ser^WA^ or Thr^WA^) that binds the Mg^2+^ ion (Fig. 6A, C). The WB motif comprises a conserved acidic residue (Asp^WB^ in F/N- and A/V-ATPases) preceded by a stretch of four hydrophobic residues. Also obligatory is the “catalytic” acidic residue (a Glu^cat^ residue in F/N- and A/V-ATPases), which serves as an immediate acceptor of a proton from the catalytic water molecule (H_2_O^cat^). While the resulting OH^−^ anion attacks the terminal phosphorus atom, the accepted proton appears to travel from Glu^cat^ to Asp^WB^, where it waits to be used in the final step of the catalytic reaction [86,87]. Fig. 6A and 6C show the WB and WA motifs, as well as the Glu^cat^ residues, in the catalytic sites of F- and A/V-ATPases, respectively. In Fig. 6A and 6C, these sites contain adenosine-5’-(β,γ-imido)triphosphate (ANP), a non-hydrolyzable ATP analog. Its binding brings the catalytic site into a “pre-transition state” [1,86,87]. The non-catalytic subunits of F- and A-ATPases lack some of these essential elements, but differently. In the non-catalytic sites of F-ATPases the Glu^cat^ is replaced by a Gln residue (Gln208 in Fig. 6B), which is unable to accept a proton from H_2_O^cat^ [1]. In addition, the Walker A motif of the P-loop contains bulky Arg and Gln residues (Arg171 and Gln172 in Fig. 6B), which prevent the insertion of the stimulating Arg residue (Arg372 of the adjacent α-subunit in Fig. 6B).

**Fig. 6.**
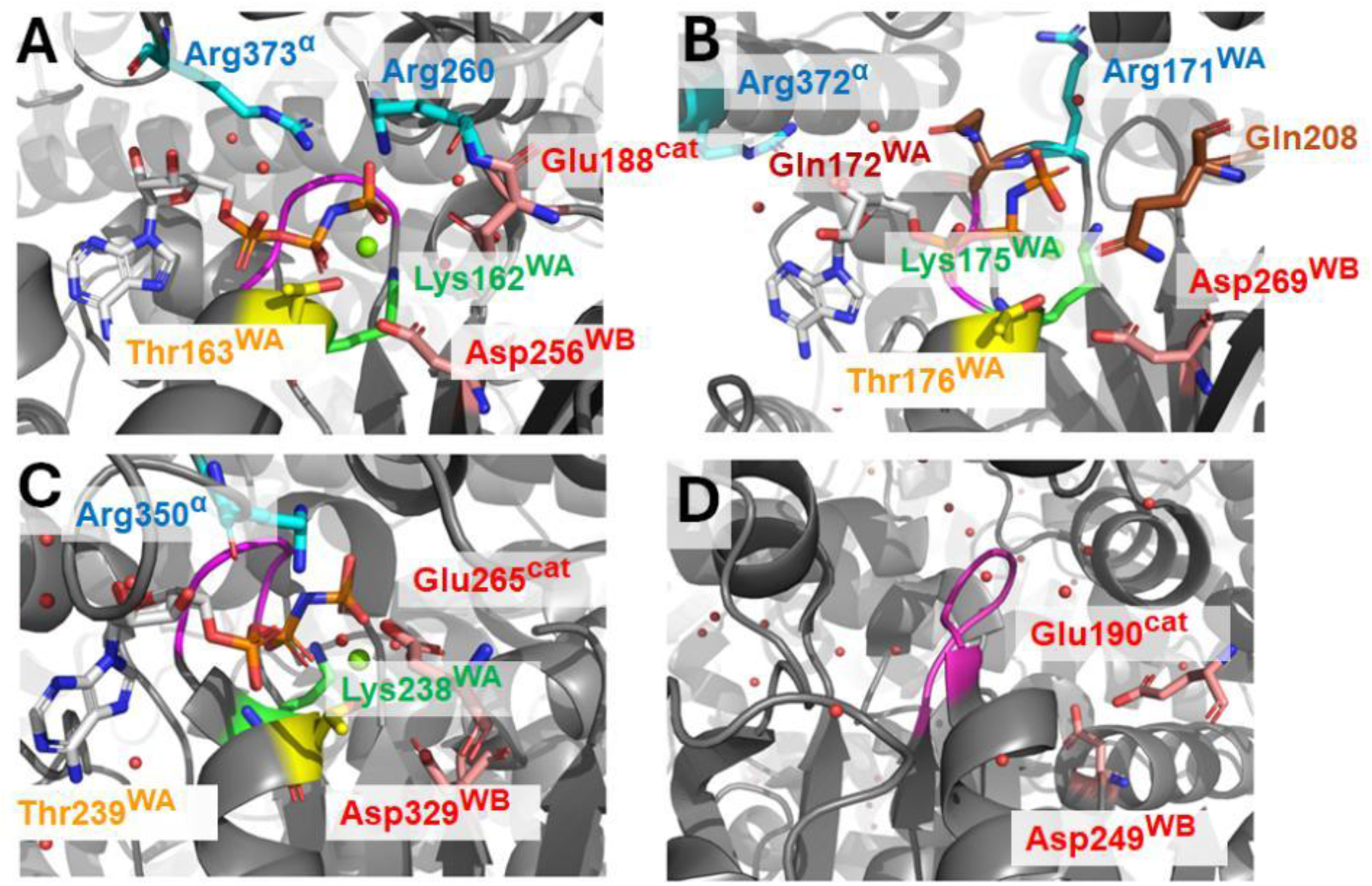
Structures of catalytic and non-catalytic sites in F- and A-ATPases. **A, B,** catalytic (**A**) and non-catalytic (**B**) sites of an F-type ATP synthase from bovine mitochondria (PDB ID 1BMF [1]. **C, D,** catalytic (**C**) and non-catalytic (**D**) sites of an A-type ATP synthase from *Enterococcus hirae* (PDB ID 3VR6 [88]). Color code: The key amino acid residues and molecules of adenosine-5’-(β,γ-imido)triphosphate (ANP, present in panels A, B, and C) are shown as balls and sticks. Color code: oxygen, red; nitrogen, dark blue; phosphorus, orange; carbon atoms are colored pink for Asp and Glu, cyan for Arg, brown for Gln, green for Lys^WA^, and yellow for Thr^WA^; the P-loop (Walker A motif) is magenta, polypeptide chains are gray. Water molecules are red spheres. Mg²⁺ ions are lime spheres.

In contrast, the non-catalytic sites of A/V ATPases lack half of the Walker A motif so that the respective protein loop is dramatically truncated preventing ATP binding (Fig. 6D). At the same time, both the Walker B motif (topped by Asp249^WB^ in Fig. 6D) and the counterpart of Glu_cat_ (Glu190 in Fig. 6D) are present.

Thus, the loss of catalytic function in F/N-ATPases was caused by the loss of Glu^cat^ and prevention of interaction with the stimulating Arg residue. In contrast, the loss of function in A/V-ATPases was caused by the major truncation of the P-loop.

If the loss of catalytic function happened twice, as we suggest, then the F- and A-ATPases likely found different solutions to mechanistically couple their catalytic sites via the non-catalytic subunits. This rationale could explain why the P-loop subunits of these ATPases differ more from each other than from the homohexameric bacterial flagellar ATPase (see Fig. 4).

The commonly held belief that the separation into catalytic and non-catalytic subunits happened only once and prior to LUCA was used to localize the root of the universal tree of life between bacteria and archaea [89]. We would like to emphasize that the here suggested scheme with two independent losses of the catalytic function does not affect this conclusion, since studies of other paralogous genes, whose duplication prior to LUCA is undisputed, have concurrently confirmed the rooting of the tree of life between bacterial and (eu)archaeal branches [42,90,91].

### 4.4. Comparison of operon structures for different rotary ATPases

A comparison of the operons of flagellar ATPases with those of F- and A-ATP synthases, as conducted on an expanded sample (Fig. 5), generally confirmed the data of Matzke et al. regarding the preservation of gene order in their common sub-operon, which comprises the genes of the (i) outer stalk, (ii) P-loop ATPase, and (iii) inner shaft [49].

Fig. 5 and Supplementary File 3 show that “flagellar” gene order is inherent in the A- and F/N-ATPases of the basal clades lying closer to the root in Fig. 4. Remarkably, flagellar ATPases have only one P-loop ATPase gene, whereas rotary ATPases have two homologous ATPase genes. Notably, in A-ATPases, the first gene encodes the catalytic subunit, while the second encodes the non-catalytic subunit. In F-ATPase, by contrast, the gene for the non-catalytic subunit comes first. This difference may be a manifestation of independent losses of catalytic functions in A- and F-ATPases, as discussed in the previous section. In A-ATPases, the first gene in the pair likely lost its function, and in F-ATPase, the second.

The “canonical” order of the F_1_ genes is δ – α – γ – β – ε (see Fig. 1B). Looking at Fig. 5, one might suggest that the object of duplication in F-ATPases was not the single gene encoding the catalytic subunit but rather two genes: one encoding the P-loop subunit (homologous to the FliI) and the other encoding the γ-subunit of inner stalk homologous to the FliJ subunit of flagella. In this case, duplication would result in pairs of α + γ and β + γ′ subunits, where γ’ denotes a paralog of the γ-subunit. We suggest that this paralog evolved into the ε-subunit. The following evidence supports this suggestion: (i) the β-ε gene pair frequently moves together across genomes, see Fig. 5 and Supplementary Figs S1 and S3. In N-ATPases, for example, they moved together to the beginning of the operon, see Fig. 5; (ii) the ε-subunit cooperates with the γ-subunit in forming the central shaft of F/N ATPases, see Fig. 1A; (iii) the ε-subunit, which is usually compactly folded, can open like a pocketknife in some organisms. Its C-terminal α-helical segment stretches then along the γ-subunit and blocks its rotation in the direction of hydrolysis [28,92]. All this may indicate that the ε-subunit originated from the paralog of the gene that coded the ancestral form of the γ-subunit.

### 4.5. Evolutionary scenario

The identified "basal" A- and F-ATPases are similar in that their operons contain fewer distinct genes than those of their predominant "typical" prokaryotic counterparts (Fig. 5 and Supplementary File 3). At the same time, it is well known that eukaryotic rotary ATPases contain even more distinct subunits [7]. Apparently, new subunits were continuously acquired over the course of evolution. Below, building on presented data, we provide an evolutionary scenario that rationalizes these acquisitions, see Fig. 7.

**Fig. 7.**
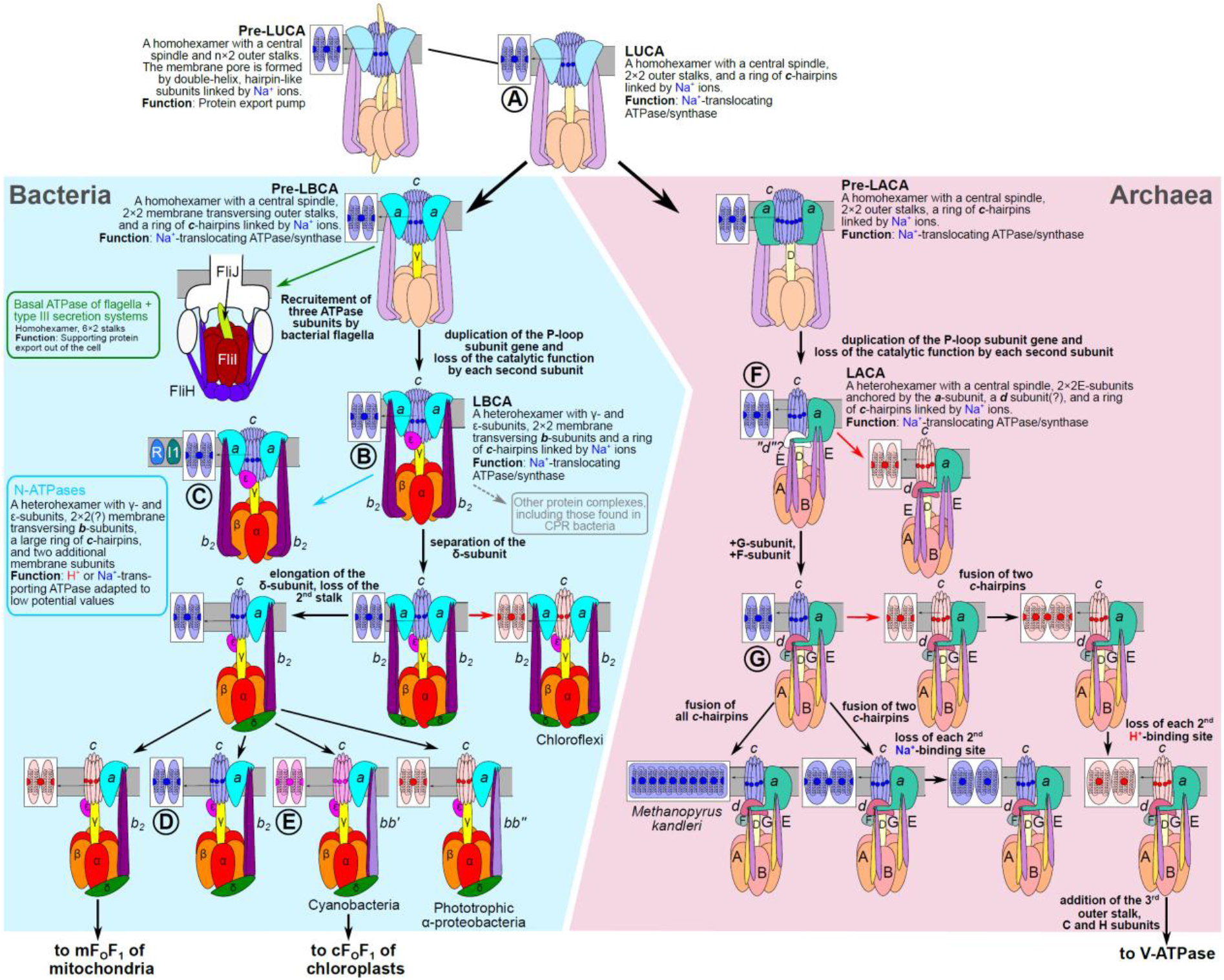
Tentative scheme of the evolution of membrane rotary ATPases. The evolution of the rotary ATPases is depicted as going from complexes with large ***c***-rings, adapted to work at low membrane potentials, to ATPases with smaller ***c***-rings, geared for high membrane potentials (see the text for details). The color code for the subunits is the same as in Fig. 1, except for the ***c***-subunits. They are colored according to their ion specificity. The Na⁺-transporting ***c***-rings are light blue, and the H⁺-transporting ***c***-rings are pink. Blue dots indicate Na⁺-binding sites, and red dots indicate H⁺-binding sites. The ***c***-ring of the cyanobacterium *Synechocystis* sp. PCC 6803 having promiscuous ion specificity [60] is shown in shades of magenta. The ATPases designated by Latin letters in squares have the following amino acid motifs in their ***c***-subunits: 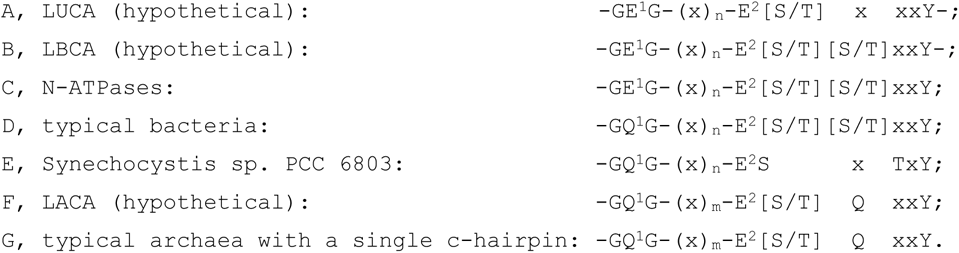

#### 4.5.1. Evolution of rotary ATPases up to LUCA

Although there is little data from the pre-LUCA era, P-loop ATPases were likely present because all of their major classes can be traced back to LUCA [93]. Interestingly, the threading of proteins through the central holes of hexameric rings is not only typical of rotary ATP synthases and flagellar ATPases belonging to the RecA/F_1_ class but also of unfoldases belonging to the AAA+ class of P-loop ATPases [94], Thus, such constructs may have been common even during the pre-LUCA era.

LUCA may have been a community of nano-sized cells that depended on sharing gene products due to their small genomes [42,95]. This exchange required membrane transporters for proteins and polynucleotides. We previously hypothesized that one of these transporters was a ring-shaped homohexamer of P-loop ATPases connected by two stalks to a membrane pore composed of hairpin-shaped subunits linked and stabilized by Na⁺ ions [37,39]. A comparison of the amino acid sequences of the c-subunits of A- and F-ATPases indicates that their hairpins arose from the duplication of a single α-helix containing a Glu residue. By linking these Glu residues, Na⁺ ions keep the membrane hairpins together, thereby strengthening the pore [96].

We previously suggested that a mutation in the pore could cause the lodging of a translocated protein, resulting in the ATP-driven rotation of the entire pore relative to the membrane subunits that anchor the complex [37]. However, the subsequently discovered homology between the central stalk of rotary ATPases and the FliJ subunit of the flagellar basal ATPase [49,84,97] suggests that it was not the transported protein that may have become stuck, but rather the central rod itself. Similarly to the FliJ subunit of flagellar ATPase, this central rod may have assisted in protein translocation (Fig. 7). In any case, such an ‘unlucky’ translocase may have given rise to ion-translocating membrane rotary ATPases [37].

The homology between subunits of A- and F-ATPases implies that the stator of LUCA’s ATPases contained a hexamer of catalytic P-loop subunits anchored in the membrane by the ancestors of the ***a***- and ***b***-subunits, see Fig. 7. The rotor part likely included a ring of hairpin-like-***c***-subunits and a coiled-coil protein, anticipated common ancestor of bacterial γ- and archaeal D-subunit [49,97].

Being unable to translocate proteins, such a rotating device would only have been preserved by evolution if it were useful. And there was a use for such a device – it may have initially helped to maintain the K^+^/Na^+^ disequilibrium over the cell membrane. The protein synthesis machinery, which emerged long before LUCA [40–42], can only function if the K^+^ level is approx. 10 times higher than that of Na⁺ [43,98]. This suggests that the first cells formed in environments with prevalence of potassium over sodium. Most likely, they may have emerged in geothermal fields overlying vapor-dominated geothermal systems, since K^+^ prevails over Na^+^ in the geothermal vapor [44,99,100]. Both freshwater and seawater contain more Na^+^ than K^+^, so cells must have developed mechanisms to maintain high levels of K^+^ in the cytoplasm in order to disperse from their places of origin or survive during rainy periods. The simplest way for it was to selectively expel Na⁺ ions. A „jammed” protein translocase with a rotating, Na^+^-stabilized ring could eventually couple the hydrolysis of ATP with transmembrane Na⁺ translocation along the interface between the “stator” and “rotor” [38,39,45]. Given the importance of the Na⁺-expelling function for the first cells, it is not surprising that several other membrane systems, which use other energy sources to pump Na^+^ out, are also traceable back to LUCA and beyond [101–104].

The evolutionarily primacy of maintaining the K^+^/Na^+^ gradient is also supported by the data of Harold and Bruns who collapsed the membrane potential in *Streptococcus faecalis* (currently named *Enterococcus hirae*) by gramicidin and other ionophores. After such treatment, the bacteria grew at the same rate on a potassium-based, nutrient-rich medium, but not on an equally nutrient-rich, but sodium-based medium [105]. Clearly, an ion-tight membrane is paramount for maintaining the K^+^/Na^+^ disequilibrium across the cell membrane but not for synthesis of ATP, which could also be produced through fermentation.

Thanks to the combined action of rotary ATPases and other Na⁺ export pumps, Δμ̅ Na⁺ must have built up at the membrane of early cells. Unlike other Na⁺ pumps, the rotary ATPase—by simply rotating in the opposite direction—could couple the discharge of this Na⁺ potential with ATP synthesis. As argued elsewhere, the interplay between the Na^+^-translocating rotary ATP synthases and other Na^+^-exporting systems may have led to the emergence of Na⁺-dependent membrane bioenergetics [38,39,45,47]. Its essence lies in harvesting and accumulating small energy quanta for ATP synthesis [45,106,107]. Indeed, the expulsion of a Na⁺ ion from the cell by a Na⁺-export pump requires only a small amount of free energy — or even none at all if the Na⁺ ions are translocated together with negatively charged “waste”, such as organic acids — common end products of fermentation. The resulting Δμ̅ Na⁺ can then drive Na⁺ ions back into the cell via ATP synthase. The ATP synthase accumulates the energy of Δμ̅ Na⁺ by converting the energy of successively conveyed Na^+^ ions into the elastic deformation of the peripheral stalk(s). When the energy stored in the deformation reaches a threshold of ∼ 60 kJ/mol, it is discharged through synthesis of an ATP molecule [7,25,45].

Yet, it is unlikely that the membranes of LUCA held the high membrane potential of 150-200 mV, which is common among present-day prokaryotes (this potential corresponds to an electric field of ∼3×10^7^ V/m over the lipid bilayer dielectric). It is believed that LUCA had primitive membranes composed of single-tail lipids [108,109]. Tighter membranes made of more complex, double-tailed lipids apparently emerged later and independently in bacteria and archaea, whose lipids are fundamentally different [110]. In addition, most reconstructions suggest that LUCA was a mesophile or moderate thermophile that lived at temperatures between 40 and 65°C [42,111]. At these temperatures, LUCA’s primitive lipid bilayer must have been permeable to protons and could have maintained only small Na^+^ gradients at best [112].

Yet, the possession of two outer stalks, as in *Chl. aurantiacus,* or three outer stalks as in genetically modified *Bacillus* PS3 [54], may have ensured the ability to synthesize ATP even at low values of ΔµNa⁺. If possessing two or three peripheral stalks, each with an ion-translocating ***a***-subunit, primordial rotary ATPases with P-loop homohexamers and large ***c***-rings of, say, 30 subunits could synthesize six ATP molecules per full rotation already at potentials as low as 40 mV (three ***a***-subunits) or 60 mV (two ***a***-subunits) by translocating 60 or 40 Na⁺ ions into the cell, respectively. By hydrolyzing six ATP molecules per full rotation, these ATPases could also effectively pump the same 40–60 Na⁺ ions out of the cell. Accordingly, Na⁺-dependent membrane bioenergetics may have been working already at the LUCA stage, being optimized for membrane potentials of ∼ 50 mV.

Since two outer stalks are found both in early branching F-ATPases, such as that of *Chl. aurantiacus* [55], and, apparently, in all A-ATPases, we envision that LUCA possessed rotary ATPases with two outer stalks, a P-loop homohexamer, and many Na^+^-binding ***c***-subunits, as shown in Fig. 7. These rotary machines were inherited by the ancestors of bacteria and archaea respectively. From this point on, the histories of bacterial and archaeal rotary ATP synthases diverge.

#### 4.5.2. On the way from LUCA

We do not know much about the transition from LUCA to bacteria and archaea, respectively. It has been hypothesized that bacteria and archaea emerged from LUCA’s descendants that were geographically isolated from each other [113].

During the time of their separate evolution, bacteria and archaea developed fundamentally different two-tail membrane lipids [110]. Apparently during the same time, the rotary ATP synthases evolved differently within lineages leading to bacteria and archaea.

##### On the way from LUCA to the Last Bacterial Common Ancestor

While the shared features of F- and A-ATPases enable the resurrection of their ancestor in LUCA, those features that are common to all F-ATPases, but not A-ATPases, enable the reconstruction of rotary F-ATPase of the Last Bacterial Common Ancestor (LBCA). Apparently, these common features were acquired along the way from LUCA to LBCA, see Fig. 7.

We believe that the genes of rotary ATP synthases, after being inherited by ancestors of bacteria, occasionally underwent duplications yielding, in particular, truncated operons carrying only cytoplasmic subunits. Such truncated operons of unknown function are still present in many bacterial genomes, see Supplementary Figure S3. We anticipate that the basal complex of bacterial flagella stems from a truncated ATPase operon, which encoded an outer stalk, a P-loop subunit, and a central rod. The products of this operon may have functioned together as an unfoldase, perhaps in conjunction with a protein translocase, predating the flagellum. All of this likely happened after the bacterial and archaeal lineages separated, since flagella are inherent to bacteria but not archaea, which use different motility machinery. The homohexameric nature of the flagellar P-loop ring suggests that the membrane rotary ATPase was also homohexameric when it was inherited by bacterial ancestors from LUCA.

A further duplication event led to the loss of the catalytic function in half of the P-loop subunits due to the loss of Glu_cat_, as discussed in Section 4.4 and shown in Fig. 7. Most likely this loss helped to save on materials and assembly logistics. As already noted, the thermodynamic efficiency of ATP synthesis is determined by the Na^+^/ATP or H^+^/ATP ratio. At a given membrane potential, only three catalytic subunits provided the same efficiency with half the number of ***c***-subunits.

As argued in Section 4.4, it looks like that the P-loop subunit duplicated together with the ancestral γ subunit in the bacterial lineage. In the α + γ gene pair, the catalytic function of the P-loop subunit was lost. In the other gene pair, the P-loop subunit β retained its catalytic function whereas the paralog of the γ subunit seemingly evolved into a smaller ε subunit, which, clinging to the γ subunit, slides under the P-loop hexamer during rotation.

Hence, the F-ATPase of LBCA already contained the catalytic and non-catalytic subunits, two outer stalks, as well as the ε subunit in support to the γ subunit, see Fig. 7.

##### Evolution of F/N ATPases after the LBCA

Initially, the F-ATPases, as discussed in Section 4.4., retained the “flagellar” order of genes coding for the outer stalk (***b***-subunit), P-loop protein (α-subunit) and inner stalk (γ-subunit); This order is conserved in the early-branching, basal F-ATPases, including the N-ATPases (Fig. 5 and Supplementary Figure S3).

The N-ATPases separated from the main trunk of F-ATPases after the LBCA (Fig. 4). N-ATPase operons are distinguished by two additional genes encoding small membrane proteins whose function remains unclear. Furthermore, they do not contain a separate gene for the δ subunit, see Fig. 7 and [48]. In addition, N-ATPases differ from other F-ATPases in gene order: the β+ε genes are at beginning of the operon in contrast to other F-ATPases where the β+ε genes follow the α + γ gene pair (Fig. 5 According to data from Osipov and colleagues, N-ATPases appear to have ∼20 ***c***-subunits [56]. As argued in Section 4.1, N-ATPases are likely to have two outer stalks.

The further evolution of F-ATPases was accompanied by a split of the outer stalk into the ***b-***and δ-subunits, which led to the loss of the “flagellar” gene order (Fig. 5). The increased tightness of bacterial membranes and higher membrane potential must have enabled a decrease in the number of *c*-subunits, as shown in Fig. 7. Yet, as long as the δ-subunit remained short, it may have enabled the accommodation of two outer stalks, as the example of *Chl. aurantialcus* shows [55]. With H^+^/ATP ratio of 6.6, the F-ATPase of *Chl. aurantialcus* is optimized for working at membrane potential of about 100 mV, which is a reasonable value for Δμ̅H^+^ in this thermophilic, phototrophic bacterium. Notably, short δ-subunits are found in several bacterial phyla, see Fig. 2 and Supplementary Figure S1.

The next change was the elongation of the δ-subunit that prevented the binding of more than one outer stalk, see Section 3.2. There are at least two possible reasons for the loss of the second stalk. First, the conversion from the two ion-conducting outer stalks to only one doubled the efficiency of ATP synthesis – provided that membranes were already tight enough to maintain membrane potential ≥ 150 mV. Second, rotation of the ***c***-ring between the two ion-conducting ***a***-subunits likely needed precise synchronization of the two ion-transfer events. The need for such synchronization may have been kinetically hampering.

The price of doubling the efficiency was the inevitable overall mechanistic destabilization of the F-ATPase complex after losing one of the two outer stalks. Obviously, the F-ATPases with only one outer stalk do operate because their γ and ε subunits together prevent the P-loop hexamer from tipping over. Mechanistically it is achieved by inelastic state of the γ-subunit [114] and tight clamping of the C-terminus of the γ-subunit by the P-loop subunits [115]. As a result, the full rotation of γ-subunit proceeds around the dihedral (Ramachandran) angles of its specific conserved glycine-containing motif and is accompanied by transient uncoiling of the peptide chain upon each turn, as shown in [115].

In general, the evolution of F-ATPases proceeded in the direction of reducing the number of ions translocated per one synthesized ATP molecule. This was made possible by bacteria becoming capable of maintaining effective Δμ̅ Na⁺ or Δμ̅ H⁺ values of around 200–250 mV — despite leaks, which increase non-linearly with rising membrane potential [116]. Specifically, the emergence of oxygen respiration, which is energetically lucrative, may have enabled lower H^+^/ATP ratios [47]. Among bacteria, Mycobacteria appear to have the lowest H^+^/ATP value of 3.0 with nine ***c***-subunits [117]. These bacteria are capable of oxygen respiration, which provides sufficient energy to maintain a high membrane potential.

Thus, as bacterial F-ATPases evolved, the number of ions translocated per one synthesized ATP molecule likely decreased from over ten to three in mycobacteria.

##### On the way from LUCA to the Last Archaeal Common Ancestor

Unfortunately, we have less structural information about A-ATPases than about F-ATPases. Thus, to date, there is no complete structure of any archaeal A-type ATP synthase. We can rely only on the two structures of A-type ATPases from the bacterium *Thermus thermophilus* [4,118]. Therefore, below we mostly refer to the sequence data and the structures of respective operons.

As with F-ATPases, the features that all A-ATPases have in common but that are absent in F-ATPases point to acquisitions made by the ancestors of A-ATPases during their evolutionary journey from LUCA to the last archaeal common ancestor (LACA)., see Fig. 7.

First, the gene of the P-loop subunit underwent duplication followed by the loss of catalytic function in one of the paralogs due to the truncation of the P-loop motif, see Section 4.4 and Fig.6. Secondly, the ATP yield was doubled – and in a more elegant manner than in F-ATPases. Specifically, the ion-translocating ***a***-subunit somehow attained extensive cytoplasmic protrusions. Then the N-termini of outer stalks began to bind to these cytoplasmic protrusions rather than embedding themselves in the membrane bilayer and binding to the transmembrane helices of the cognate ***a***-subunit there. Finally, two outer stalks ended up bound to the same single ion-translocating ***a***-subunit as seen in Fig. 1A.

To stabilize the whole construct, the archaeal ***a***-subunit attained four additional intramembrane helices, as compared to the bacterial ***a***-subunits. These four helices are additionally anchoring the stator in the lipid bilayer. Obviously, with two outer stalks and a fester membrane-anchored ***a***-subunit, the A-ATP synthases are mechanically more stable than F-ATPases having only one outer stalk [21].

##### Evolution of A-ATPases after the LACA

Our data are ambiguous concerning two other acquisitions by A-ATPases, namely the conical ***d***-subunit serving as a socket for the central stalk, and the F-subunit, which clings to this stalk. We failed to identify their genes in some of the operons of “basal” A-ATPases. Hence, the acquisition of these subunits may have occurred during the evolutionary transition from basal A-ATPases (group of clades 2 and clade 3) to canonical ones (Clade 1), see Fig. 3 and Section 3.2. The G-subunit, which together with the homologous E-subunit forms the outer stalk of most A-ATPases, is also absent from the operons of basal A-ATPases. It is likely that the G-subunit emerged from a duplication of the E-subunit gene upon the transition from basal to canonical A-ATPases.

The present-day A-ATPases have more diverse ***c***-subunits than F-ATPases [18,21]. The duplications of ***c***-subunits are much more widespread among A-ATPases than in F-ATPases. It is believed that the ancestral A-ATPase had hairpin-like ***c***-subunits with only two transmembrane helices [18,21]. Such hairpins are found in some nanoarchaea and methanogens, which are believed to be early-branching archaea. Yet many methanogens have duplicated ***c***-subunits consisting of two nearly identical hairpins, each providing a binding site for one Na⁺ or H⁺ ion [18,21,46]. However, the ***c***-subunits of many archaea, such as Pyrococci, as well as A-ATPases found in bacteria, have four transmembrane helices that provide a binding site for only one Na⁺ ion (e. g. in *Enterococcus hirae* (see Fig. 1C and [18,20,21]) or one proton (e. g. in *Thermus thermophilus,* see Fig. 1A and [4,118]). Thus, A-ATPases can modulate the Na^+^/ATP or H^+^/ATP ratio not only by varying the number of ***c***-subunits, but also by varying the number of ions bound per one ***c***-hairpin. While two- and four-helical ***c***-subunits are most widespread in archaea [18,21], the number of hairpins in archaeal ***c***-subunits can vary widely. Ultimately, *Methanopyrus kandleri* carries a single ***c***-subunit gene coding for 1021 amino acids supposedly forming 13 hairpins [119].

In fact, the structural difference between the single-hairpin and double-hairpin subunits is not large; they can even be combined in the same oligomeric ring as in acetogenic bacterium *Acetobacterium woodii*, where the ***c***-ring contains nine single-hairpin subunits typical for F-type ATPases and one four-helical ***c***-subunit typical for A-type ATPases [120].

Due to the scarcity of structural information, data on the number of ions translocated by A-ATPases per one ATP are also limited. The highest efficiency of A-ATPases is compatible with that of F-ATPases; the lowest number of ions needed to synthesize ATP, namely 3.3, can be deduced from the structure of *Enterococcus hirae **c***-ring having 10 ***c***-subunits, see Fig. 1C and [20]. Yet, it cannot be excluded that A-ATPases of some archaea with a very tight energy budget may need to translocate more than 13 ions per full turnover.

#### 4.5.3. Evolution of ion-specificity of prokaryotic rotary ATPases

As Figures 2-4 and Supplementary Figures 1-3 show, Na⁺-binding sites in ***c***-subunits are widespread in ATPases that branch early in phylogenomic trees. These include F-ATPases found in Thermotogota and several other phyla, N-ATPases found in various lineages (see Fig. 2 and Supplementary Figure S1), as well as A-ATPases of group of clades 2 and clade 3 (see Fig. 3 and Supplementary Figure S2).

As discussed elsewhere, the binding of Na^+^ by a ***c***-ring in the middle of hydrophobic membrane requires at least six coordinating ligands, one of which is a negatively charged Glu that compensates for the charge of the Na^+^ ion [47,121]. In contrast, the minimal requirement for proton translocation is just a Glu or Asp residue in the middle of the second helix (E^2^ or D^2^, see Section 2.3); this Asp or Glu becomes electrostatically neutral upon its protonation and can enter the lipid bilayer upon rotation of the ***c***-ring. Therefore, Na^+^-ATPases can translocate protons in the absence of Na^+^ ions, but not the other way around [2].

In Fig. 7, we indicated, where appropriate, the Na^+^-binding motifs of ***c***-subunits. The motif suggested for LUCA contains Na^+^-binding side chains common for bacteria and archaea, including E^1^ and E^2^, as well as a conserved Tyr in the E^2^+5 position, which stabilizes E^2^, see Fig. 1C and [47].

In many early branching F-ATPases the ***c***-subunit has a Glu residue in the middle of the first helix (E^1^). This “primordial” motif is still present in many N-ATPases, including those few found in archaea [48,52,53,122]. Also, E_1_ and E_2_ are found among fusobacteria, such as *Fusobacterium nucleatum*, see Fig. 1C [24,48]. In the case of the archaeon *Methanosarcina mazei*, which contains both a H^+^-translocating A-ATPase and an N-ATPase with the supposedly primordial Na^+^-binding motif, the ability of its N-type ATPase to couple Na^+^ transport with ATP synthesis at pH 6.9 has been shown experimentally [123].

Remarkably, in many Na^+^-translocating ATPases of bacteria and archaea, the E^1^ residue got replaced by Gln (Q^1^). It is tempting to speculate that the substitution of noncharged Gln for E^1^ decreased the probability of the ***c***-ring destabilization in case of eventual deprotonation of this glutamate. While E^1^ can provide two oxygen atoms to coordinate a Na^+^ ion, Gln can provide only one. Perhaps therefore, the replacement of E^1^ by Gln seems to parallel the acquisition of an additional Na^+^ ligand in the E^2^+2 position. In bacteria, this additional ligand is a Ser/Thr that coordinates the Na^+^ ion via a water molecule see Fig. 1C and [121]. In the cognate position, archaea have Gln whose longer side chain coordinates Na^+^ directly, see Fig. 1C and [121].

Figure 7 incorporates the recently demonstrated ion promiscuity of the sole ATP synthase found in the cyanobacterium *Synechocystis* sp. PCC 6803. Forchhammer and his colleagues showed that this F-ATPase, having Q^1^ and all “canonic” Na^+^-binding side chains except the one in the E^2^+2 position (see Fig. 7), translocates protons at acidic and neutral pH, but switches to the Na^+^ ions at pH > 8.0 [60]. Such an adaptation is extremely useful for phototrophic cyanobacteria: the ATP synthases in membranes of intracellular thylakoids harvest protons from their lumens having pH of ∼6.0 at light [124], whereas the same ATP synthases in the outer membrane, harvesting ions from the environment with a pH of ∼8.0 (e.g., seawater), can synthesize ATP using the Na⁺ gradient present over the outer membrane (e.g. in the dark). Apparently, the absence of the Na^+^-binding side chain in the E^2^+2 position – or its replacement by less fixed water molecule – weakens the binding of a Na⁺ ion, so that it can be ousted by a proton at pH <8.0. Intriguingly, the promiscuous motif is present in about half of genomes of oxyphototrophic cyanobacteria, whereas the other half has a similar motif, but with two Na^+^-binding side chains (E^2^+1 and E^2^+2) missing. It would be interesting to determine experimentally whether the F-ATPases of these cyanobacteria are incapable of Na⁺ translocation or if they switch to it at even higher pH values (>9.0).

Remarkably, *Methanosarcina acetivorans*, like the previously mentioned *Methanosarcina mazei*, has both an A-ATPase and an N-ATPase with the "primordial" Na⁺-binding motif having E^1^ and E^2^ [48]. In these archaea, the ATP-driven Na^+^ translocation was observed at pH 7.0, but not at pH 5.0 [125]. These data may indicate that the primordial Na^+^-binding motif, which we attributed to LUCA, might switch from binding of Na^+^ ions to protons at acidic pH < 5.5. If so, then the above-mentioned acquisitions of additional Na^+^-binding side chains in the E^2^+2 position may have been also driven by the need to increase the specificity of Na^+^ binding.

Otherwise, Fig. 7 depicts the multiple transitions from the Na^+^ translocation to the H^+^ translocation following the loss of Na^+^-ligands. As discussed elsewhere, the proton-dependent bioenergetics became more advantageous with appearance of atmospheric oxygen – given that the cell membranes were already proton-tight [39,45,47,112]. These transitions to proton-dependent bioenergetics occurred independently in distinct prokaryotic lineages, as justified by the sequence analysis in our previous publications [38,46].

### 4.6. Rotary membrane ATPases evolved to increase the net ATP production

Apparently, the evolution of rotary ATP synthases was accompanied by an increase in net ATP production, definable as the difference between the amount of ATP synthesized and hydrolyzed by these enzyme complexes. Most of the ATP produced (about 75%) is used for the biosynthesis of proteins and polynucleotides [126]. The maximization of available ATP for these cellular processes has been achieved through several complementary means, which will be discussed below.

#### 4.6.1. Optimizing the ATP distribution and utilization

One way to increase the supply of ATP for biopolymer synthesis is to optimize its use in housekeeping processes. This optimization can be traced for systems maintaining the K^+^/Na^+^ disequilibria over the cell membrane. These systems are essential consumers of both ATP and membrane potential (see Section 4.5.1).

Specifically, some ancestors of methanogenic archaea developed membrane Na^+^/K^+^-ATPases [127]. Unrelated to rotary ATPases, Na^+^/K^+^-ATPases transports three sodium (Na^+^) ions out of the cell and two potassium (K^+^) ions into the cell for every ATP molecule hydrolyzed [128]. Therefore, Na^+^/K^+^-ATPases are nearly twice as efficient as enzymes of membrane bioenergetics at maintaining the K^+^/Na^+^ disequilibrium.

The further optimization of ATP consumption is coupled with the emergence of eukaryotes. They most likely evolved from a syntrophic consortium containing archaea and bacteria, whose members merged to form a new organism [129,130]. The information-processing machinery of this new organism, including translation, came from the ancestors of Asgard archaea. The largest proportion of bacterial proteins originated from ancient myxobacteria [130]. The rest of the genes came from a variety of organisms, supposedly via horizontal gene transfer [130]. Specifically, the K^+^/Na^+^ disequilibrium in eukaryotes is maintained by Na^+^/K^+^-ATPases alike those in methanogens [127], although it is unclear yet how their genes made it from methanogenic archaea into the eukaryotic genome.

In a separate event, α-proteobacteria were acquired as endosymbionts and developed into mitochondria. Additionally, eukaryotic lineages that already possessed mitochondria acquired cyanobacteria as another type of endosymbiont that evolved into chloroplasts [131]. Mitochondria and chloroplasts reside within eukaryotic cells, where the K^+^/Na^+^ ratio is maintained at ∼10 thanks to the activity of Na^+^/K^+^-ATPases in the cell envelope. Therefore, these organelles spend neither membrane potential nor ATP to keep K⁺ ions in and Na⁺ ions out. Consequently, the membrane potential of the mitochondrial inner membrane is primarily used for ATP synthesis. Nearly all the ATP produced by mitochondria and chloroplasts is exported into the cell cytoplasm. Interestingly, part of it is used to power the Na^+^/K^+^-ATPases in the cell membrane. Thus, the F-ATPases still care for the K^+^/Na^+^ disequilibrium at the eukaryotic cell membrane, albeit indirectly.

In eukaryotes, ATP synthases are not localized in the outer membrane as they are in most prokaryotes. Instead, they are spread throughout the cytoplasm, carried by mitochondria and chloroplasts. This transition from 2D to 3D enables the keeping the ATP level high throughout large eukaryotic cells. Furthermore, ATP synthesis in mitochondria and chloroplasts is driven by powerful energy processes: oxygen respiration and oxygenic photosynthesis, respectively. Therefore, eukaryotic cells can have more ATP at their disposal than most prokaryotic cells [132].

#### 4.6.2. Inhibition of unnecessary ATP hydrolysis under low-energy conditions

The ATP to ADP ratio in the cytoplasm of both prokaryotes and eukaryotes is approximately 30, which is not particularly favorable for ATP synthesis. Yet, the prevalence of ATP over ADP seems to be needed to keep the ATP-consuming reactions running smoothly. Accordingly, at an ATP/ADP ratio of ∼30, the ATP hydrolysis is thermodynamically more favorable than the ATP synthesis. Under physiological conditions, rotary ATPases are prevented from hydrolyzing ATP only by backpressure from the membrane potential [12,13]. However, under low-energy conditions (e.g., food limitation), the membrane potential can drop, enabling ATP hydrolysis.

To prevent unnecessary ATP hydrolysis under low-energy conditions, rotary ATPases use several mechanisms, namely:

**a) Inhibition by the ADP-Mg²⁺ complex**: When this complex binds tightly to a catalytic site in the absence of inorganic phosphate, the catalytic pocket closes tightly around the bound complex. This locks the rotary ATPase into a stable, non-functional state, which halts rotation and catalytic activity completely. The probability of this “locking” increases when the level of ATP and/or phosphate in the cell comes down, and the level of ADP increases. The “ADP-Mg²⁺-blocked” ATPase can be activated by the recovery of a high membrane potential and/or physiological levels of ATP and phosphate [92,133–135]. This mechanism is common to F- and A-ATPases [92,133–135], so it was likely present already in LUCA.
**b) Inhibition by specific proteins:** complementary to ADP-Mg²⁺-inhibition, several mechanisms that are inherent to groups of organisms have been described, namely: *Blockage of ATP hydrolysis by the ε-subunit*: The closed conformation of the *ε-*subunit is stabilized by a bound regulatory ATP molecule. When the ATP level in the cell goes down, this ATP molecule detaches, and the epsilon subunit “opens” as a pocketknife. It pushes its C-terminal α-helix, along the coil-coiled γ-subunit, deeply into the P-loop hexamer blocking its rotation [28]. *Blockage of ATP hydrolysis by the ζ subunit in α-proteobacteria: I*n some *α-*proteobacteria (such as *Paracoccus denitrificans*) the rotation in the direction of ATP hydrolysis can be additionally blocked by a separate ζ subunit. It inserts itself into the αβ catalytic interface of the F₁ domain to retard hydrolysis [136]. This α-bacterial ζ subunit was found to be an evolutionary predecessor of the well-studied *inhibitor factor 1 (IF1)* that blocks the unwanted ATP hydrolysis by mitochondrial F-ATPase (mF1F_O_). In mitochondria, IF1 binds to mFoF_1_ when ΔµH^+^ drops and, consequently, the pH in the matrix decreases [137]. *Inhibition of ATP hydrolysis in cyanobacteria by a small protein Atpθ.* This protein, widely conserved in cyanobacteria, becomes expressed under low-energy conditions, such as darkness, and inhibits ATP hydrolysis in a dose-dependent manner [138].
**с) Blocking the ATP hydrolysis by a disulfide bridge**. The ATP synthases of plant chloroplasts (cF_O_F_1_) apparently lack the *Atpθ* subunit. But they have two cysteine residues in their γ-subunit which make a disulfide bond under low-energy conditions (e.g. in the night) and physically lock the rotor to prevent ATP consumption. In the light, thioredoxin reduces this disulfide bridge and cleaves it, using electrons coming from the working Photosystem I [139].
**d) Inhibition of hydrolysis via detachment of the V_1_ part in eukaryotic V_O_V_1_ ATPases.** In eukaryotic V_O_V_1_ ATPases, the unwanted ATP hydrolysis is prevented by the controlled reversible detachment of the entire V_1_ part, which becomes inactive after detached. The V_O_ part, in turn, loses its proton-conducting capacity once separated from V_1_ [140–142].

#### 4.6.3. Modulating the thermodynamics of ATP synthesis to increase the ATP yield

In the abovementioned cases, the level of ATP is maintained by simply preventing its hydrolysis by rotary ATPases. However, the net ATP production can also be enhanced by regulating the thermodynamics of ATP synthesis, as discussed below.

**a) Increasing the net ATP production through decreasing the Na^+^/ATP and H^+^/ATP ratios**: Apparently, the above considered transition from the “primitive” F-ATPases (those of basal clades and N-ATPases) to their “canonical” versions was accompanied by a decrease in the number of ions translocated for synthesis of one ATP molecule. Thus far, the N-ATPase from *Burkholderia pseudomallei* has the largest experimentally determined number of ***c***-subunits — 17 [70]. If this ATP synthase has only one peripheral stalk, then the ion/ATP ratio is 5.66. However, if the above suggestion of two peripheral stalks in N-ATPases is correct, then 34 ions would be translocated per one turnover. This would give an ion/ATP ratio of 11.3. For the F-ATP synthase of *Chl. aurantiacus* with two outer stalks, the H^+^/ATP ratio is 6.6 [55]. Among the F-ATPases with one peripheral stalk, the highest H^+^/ATP ratio, 5.0, can be inferred for the cyanobacterium *Arthrospira platensis* [143] and the lowest ratio, 3.0, for mycobacteria [117]. For the A-ATP synthases, because of scarcity of data, the inferred variability in the cation/ATP ratio is modest – from 4.3 in in *Methanopyrus kandleri* [119] to 3.3 in *Enterococcus hirae* [20]. However, it cannot be excluded that the ratio could be > 4.3 in some archaea on a low energy budget. So far, all ***c***-rings in available V-ATPase structures consist of ten subunits, although these *c*-subunits may have somewhat different amino acid sequences [30,144]. The chloroplast ATP synthase (cF_O_F_1_) of spinach has 14 subunits, same as in some cyanobacteria [145]. Remarkably, the number of ***c***-subunits is 13 in the cF_O_F_1_ of green algae *Chlamydomonas reinhardtii* [146]. In all resolved structures of mitochondrial F-ATPases (mF_O_F_1_), except that of animals, the number of ***c***-subunits is ten [7,11]. The mF_O_F_1_ of animals (Metazoa) have only eight ***c***-subunits [6,147]. Remarkably, the α-proteobacterium *Paracoccus denitrificans*, which is closely related to bacterial ancestors of mitochondria [148], has 12 ***c***-subunits [3]. Hence, the H^+^/ATP ratio decreases from bacteria to animal mitochondria. In animal mitochondria, having only eight ***c***-subunits [6,147], thermodynamics is most favorable and proton gradient is used in a most sparing way: ATP synthases need to translocate just 2.7 protons to produce one molecule of ATP, compared to 4 protons translocated by the ATP synthase of *P. denitrificans*.
**b) Decreasing the ATP/ADP ratio within the mitochondria**. The mitochondrial matrix, where ATP is synthesized, is separated from the cell cytoplasm by the two mitochondrial membranes. This feature allows keeping the ATP to ADP ratio in the mitochondrial matrix lower than in the cytoplasm. Indeed, the free energy of ATP synthesis (ΔG_ATP_) in the cytoplasm is about −60 kJ/mol (at ATP/ADP ratio ∼ 30) [12,149,150]. Then the threshold Δμ̅H^+^ for ATP synthesis in animal mitochondria with only eight ***c***-subunits can be estimated to be ∼ 225 mV. This value is high enough to slow the respiratory chain, causing the generation of reactive oxygen species (ROS) [12]. To avoid this, eukaryotic cells maintain a low ATP to ADP ration in mitochondrial matrix [12,150]. The membrane potential pushes synthesized ATP molecules, carrying four negative charges, out of the mitochondria through adenine-nucleotide transporters (ANT), the most abundant proteins in the inner mitochondrial membrane [12,150]. Consequently, the ATP to ADP ratio inside the mitochondria remains < 1.0 which corresponds to ΔGATP of > –45 kJ/mol [150] and enables ATP synthesis already at Δμ̅H^+^ just over 170 mV. In animals, this mechanism seems to be further enhanced by the kinases present in the intramembrane space and on the outer mitochondrial membrane. They intercept the ATP molecules leaving the ANTs and use them to phosphorylate their cognate substrates. The product ADP molecules directly return into the mitochondrial matrix via ANTs. This mechanism further ensures the low ATP to ADP ratio within animal mitochondria and enables synthesis of ATP at lower Δμ̅H^+^ levels *in vivo* [151–161].

All the manipulations of the ATP/ADP ratio described above are usually considered as means of avoiding the production of ROS by the respiratory chain [162,163]. However, any increase in the thermodynamic favorability of ATP synthesis results in the retardation of ATP hydrolysis, which directly increases net ATP production.

#### 4.6.4. Increasing the net ATP production by improving the robustness of ATP synthases

Net ATP production levels are also directly related to the mechanical stability of rotary ATP synthases. They are not very robust since the F_1_, A_1_, and V_1_ parts are only connected to the respective F_O_, A_O_, and V_O_ parts via hydrogen bonds and Mg²⁺-mediated salt bridges; no covalent bonds are involved. For instance, detaching the F_1_ part from the F_O_ part requires only the addition of a Mg²⁺ chelator and shaking of the probe. In V_O_V1 ATPases, such a detachment happens even physiologically, in response to a drop in cellular ATP level as noted above [140–142]. Consequently, an ATP synthase can be eventually broken down by a thermal fluctuation on a background of up to ∼60 kJ/mol energy transiently stored in elastic deformation of peripheral stalk(s). Since the loss of functional ATP-synthases inevitably leads to a decrease in the net ATP production, their mechanical reinforcement increases the ATP yield.

The F-ATP synthases having only one peripheral stalk are particularly vulnerable. The ATP synthase of cyanobacterium *Thermosynechococcus elongatus* was shown to remain highly active for just four hours and then decay [164]. During this time, one ATP synthase can synthesize 10^5^-10^6^ ATP molecules [139,164,165], which is compatible with the number of ATP molecules needed to synthesize an ATP synthase itself (1.4 x 10^5^) [166].

To increase the robustness of rotary ATPases, organisms use different means, as considered below:

**a) Mechanistic stabilization of membrane rotary ATPases by additional subunits:** In the case of F-ATPases, the elongation of the δ-subunit (see Sections 3.1. and 4.1.) strengthened the connection between the peripheral stalk and the P-loop hexamer. Furthermore, the ***b***-subunit underwent independent duplications in some bacteria, allowing the two ***b***-subunits (***b*** and ***b’***) to evolve separately and adapt to their different binding patches on the P-loop hexamer, see Fig. 7 and [167]. The A-ATPases also duplicated the gene of their peripheral stalk subunit, yielding two different subunits E und G, and additionally reinforced their ***a***-subunit, see Section 4.5.2 and [49,168]. Yet, only with emergence of eukaryotes, the reinforcement of rotary ATPases got into full swing. Obviously, eukaryotes have a less tight energy budget than prokaryotes. Hence, eukaryotic F- and V-ATPases can afford to possess more distinct subunits than their prokaryotic counterparts. The V-ATPases, structures of which have been resolved so far, contain two large eukaryote-specific subunits of the stator, C and H, see Fig. 1. Subunit C serves as a console for the third outer stalk, whereas both subunits are involved in additional stabilization of the whole complex [169]. The mitochondrial FOF_1_-ATPases (mF_O_F_1_) contain many subunits absent in prokaryotes. Mostly they are involved in strengthening the outer stalk and formation of mF_O_F_1_-dimers. Since these additional subunits are different in different groups of eukaryotes [7,169], they may have emerged independently in different eukaryotic lineages. All known mF_O_F_1_ structures reveal dimers or higher order multimers. Dimerization is typically viewed as a means of folding the inner mitochondrial membrane into multiple cristae, thereby increasing the energy-converting area. For example, in animals and fungi (Opisthokonta), dimers have an angle of approximately 86° between the central stalks of the two mF_O_F_1_. These dimers form long rows that supposedly impose high local membrane curvature and form mitochondrial cristae [7]. However, in ciliates (belonging to the SAR group), the angle between the two central mF_O_F_1_ stalks varies from 0° in *Paramecium multimicronucleatum* and *Tetrahymena thermophila* to 19° in *Toxoplasma gondii*, and mF_O_F_1_ dimers tend to form higher-order structures, such as dimers of dimers or hexamers of dimers [7]. These nearly flat, multimeric mFOF_1_ complexes are unlikely to impose significant membrane curvature. At the same time, in all structurally resolved mF_O_F_1_ dimers so far, their peripheral stalks interact and stabilize each other through additional subunits [7]. It is tempting to suggest that, in eukaryotes, the dimerization primarily mechanically stabilizes the mFOF_1_ complexes and only secondarily contributes to shaping the inner mitochondrial membrane. In the dimeric mF_O_F_1_ of plants, exemplified by the ATP synthase of a unicellular green algae *Polytomella* sp., the peripheral stalks are also very massive because of presence of many additional plant-specific subunits [7]. In contrast to plant mF_O_F_1_, the ATP synthases of plant chloroplast (cF_O_F_1_) have retained almost the same subunit content as cyanobacterial ATP synthase [7]. It is unlikely that this conservativity is due to the energy shortage in chloroplasts. Rather, it reflects the extreme conservation of the entire chloroplast proteome. This set of proteins, which is resistant to the oxygen it produces, emerged around 2.5 – 3.0 billion years ago [170,171] and has remained virtually unchanged in diverse cyanobacterial lineages and chloroplasts ever since. Remarkably, there is evidence of cF_O_F_1_ dimerization under physiological conditions, although the shape of these dimers remain controversial [10,172,173].
**b) Borrowing more (thermo)stable rotary ATPases via lateral gene transfer**: Last but not least: Since the A-ATPases are more (thermo)stable than F-ATPases [21], the obvious solution for bacteria to secure their net ATP production was to borrow a more robust A-ATPase from archaea via lateral gene transfer. As the data in Fig. 3 and Supplementary Fig. S2 show, such borrowing happened at least two times, each time being followed by the spread of A-ATPases among bacteria, particularly thermophilic ones. We believe that the higher (thermo)stability of A-ATPases explains why they are much more common in bacteria than F-ATPases are in archaea, see [52,53].

## Acknowledgements

Very useful discussions with Drs. D.A. Cherepanov, W. Junge, B.A. Feniouk, M.Y. Galperin, E.V. Koonin, K.S. Makarova, V.P. Skulachev, and Y.I. Wolf, are greatly appreciated. This work was supported by the Russian Science Foundation, project No. 25-24-00757 to DVD, https://rscf.ru/project/25-24-00757/

