## Supplementary material for "Evolution of membrane rotary ATP synthases: A trade-off between the net productivity and mechanistic stability?": Captions to Supplementary Figures S1, S2, and S3

**Captions to the Supplementary Figures for the manuscript:**

***Figure S1*.** **Phylogenetic tree for the β-subunits of F-type ATPases (COG0055)**. This tree is a "rectangular" version of the tree shown in ***Fig. 2*** of the main text. Details on the tree construction are given in the caption to ***Fig. 2.*** The names of leaves are colored according to the taxonomy of the respective species, the color code is shown in the box at the top of the figure.

Blue bars show the predicted number of ***c***-ring subunits according to (Osipov *et al.*, 2025 Organisms with experimentally established numbers of C-ring subunits are given in a larger font.

Green bars show the lengths of the δ-subunits. In cases of fusion between the δ-subunit and either the α- or β-subunit, the green bars show the length of the fused protein segment not covered by the respective α- or β-subunit HMM profile.

Circles at the ends of branches show the prediction of the ion specificity of the ***c***-subunits. Red circles indicate $H^+$ specificity, blue circles indicate $Na^+$ specificity, and purple circles indicate nearly complete $Na^+$-binding sites with one missing residue.

Genomic neighborhoods are visualized using the pyCOGNAT program (https://github.com/udavdasha/pyCOGNAT). The length of the arrows is proportional to gene lengths, and the colored part of the arrows is proportional to the size of the corresponding profile HMM hit. The arrow tips are not colored. The genomic neighborhood is limited to ten protein-coding genes, counting from the β-subunit of the complex. We used a set of profile HMMs (Dibrova & Rykov, 2026) to assign protein regions to Clusters of Orthologous Groups of Proteins (COGs, Galperin et al., 2025) with the hmmscan program from the HMMer package (http://hmmer.org/). The E-value threshold was set to 0.01. If two domain hits overlap by more than 5% of the length of the longer hit, the hit with the higher e-value is filtered out. Subunits of ATP synthase complexes and their predicted assembly factors are colored according to the scheme at the top of the figure. Proteins with hits from other profile HMMs are colored gray, while proteins with no COG profile HMM hits are left uncolored.

***Figure S2***. **Phylogenetic tree for catalytic A-subunits of rotary membrane A-ATPases (COG1155)**. This tree is a “rectangular” version of the tree shown in ***Fig. 3*** of the main text. Details on the tree construction are given in the caption to ***Fig. 3***. The names of leaves are colored according to the taxonomy of the respective species. The color code for taxonomy is shown in the box at the top of the figure.

Circles at the ends of branches show the prediction of the ion specificity of the ***c***-subunits. Red circles indicate $H^+$ specificity, blue circles indicate $Na^+$ specificity, and purple circles indicate nearly complete $Na^+$-binding sites with one missing residue.

Genomic neighborhoods are visualized as in ***Fig. S1*** and are colored according to the scheme at the top of the figure.

***Figure S3.*** **Phylogenetic tree of the catalytic β-subunits of bacterial F-ATPases (COG0055), the catalytic A-subunits of archaeal A-ATPases (COG1155), and the homologous flagellar ATPase FliI (COG1157)**. This tree is a “rectangular” version of the tree shown in ***Fig. 4*** of the main text. Details on the tree construction are given in the caption to ***Fig. 4***. The branches and names of leaves are colored according to the taxonomy of the respective species (the color code is the same as for ***Fig. S1*** and ***Fig. S2***).

Circles at the ends of branches show the prediction of the ion specificity of the ***c***-subunits. Red circles indicate $H^+$ specificity, blue circles indicate $Na^+$ specificity, and purple circles indicate nearly complete $Na^+$-binding sites with one missing residue.

Genomic neighborhoods are visualized as in ***Fig. S1*** except that genomic neighborhood is limited to 20 protein-coding genes colored according to the scheme at the top of the figure.
