## Supplementary figures and images for "Evolution of membrane rotary ATP synthases: A trade-off between the net productivity and mechanistic stability?"

### Supplementary Figure S1

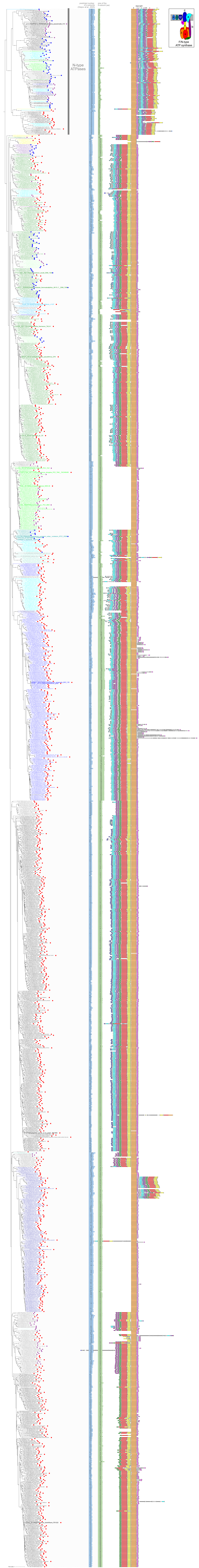

### Supplementary Figure S2

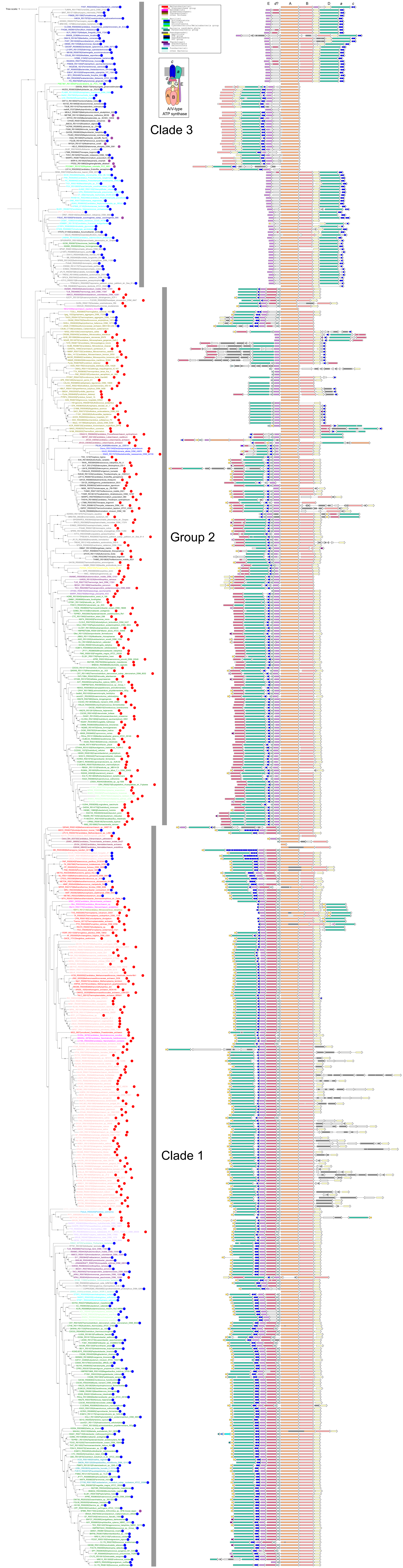

### Supplementary Figure S3

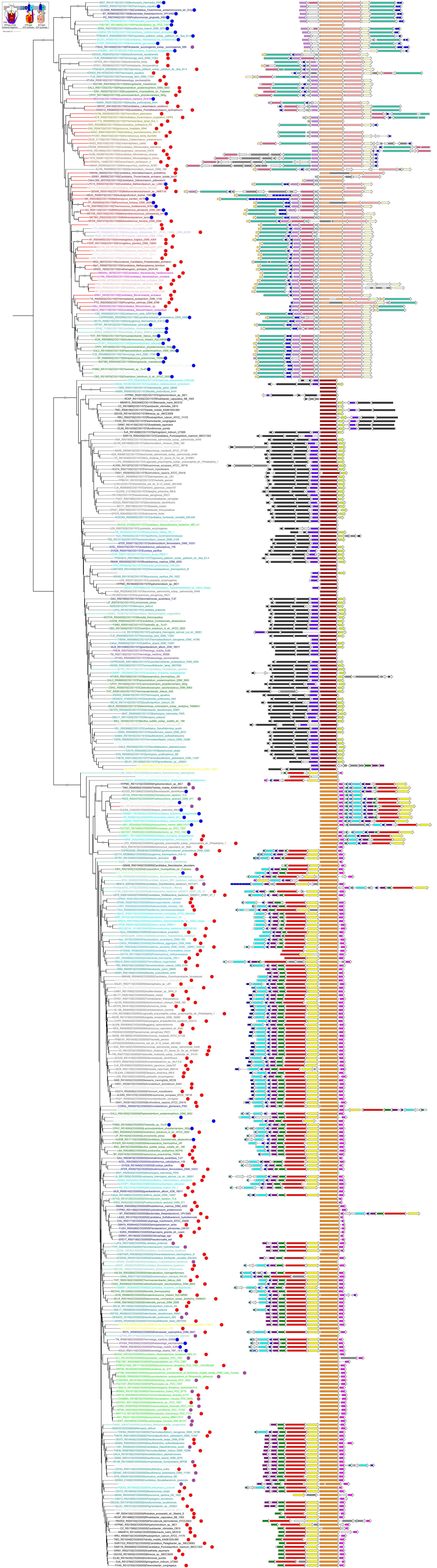
